# Cdk/Cyclin activity helps set mitotic centrosome size by influencing the centrosome growth rate and growth period

**DOI:** 10.1101/2023.04.25.538283

**Authors:** Siu-Shing Wong, Alan Wainman, Saroj Saurya, Jordan W. Raff

## Abstract

Mitotic centrosomes assemble when centrioles recruit large amounts of pericentriolar material (PCM) around themselves in preparation for cell division. How the mitotic PCM grows to the correct size is unclear. In *Drosophila* syncytial embryos, thousands of mitotic centrosomes assemble in a common cytoplasm as the embryo proceeds through 13 rounds of near-synchronous nuclear division. During nuclear cycles (NCs) 11-13 these divisions gradually slow, and we find that mitotic centrosomes respond by reciprocally slowing their growth rate and increasing their growth period so that they grow to a consistent size at each cycle. This size homeostasis is enforced, at least in part, by the Cdk/Cyclin cell cycle oscillator (CCO). Moderate levels of CCO activity appear to initially promote centrosome growth by stimulating Polo/PLK1 recruitment to centrosomes, while higher levels of activity subsequently inhibit centrosome growth by phosphorylating centrosome proteins to decrease their centrosomal recruitment and/or maintenance as the embryos enter mitosis. Thus, the CCO initially promotes, and subsequently restricts, mitotic centrosome growth to help ensure that centrosomes grow to a consistent size.

## Introduction

Centrosomes are important organisers of the cell that are formed when mother centrioles recruit pericentriolar material (PCM) around themselves (Conduit *et al*, 2015; Bornens, 2021; Vasquez-Limeta & Loncarek, 2021; Lee *et al*, 2021; Woodruff, 2021). The PCM consists of several hundred proteins (Alves-Cruzeiro *et al*, 2013), including many that help nucleate and organise microtubules (MTs), as well as many signalling molecules, cell cycle regulators, and checkpoint proteins—allowing the centrosome to function as both a major MT organising centre and an important coordination centre in many eukaryotic cells (Arquint *et al*, 2014; Chavali *et al*, 2014).

In interphase, the centrosomes in most cells organise relatively little PCM, but in virtually all cells with centrosomes there is a dramatic increase in PCM recruitment as cells prepare to enter mitosis—a process termed centrosome maturation (Palazzo *et al*, 2000; Conduit *et al*, 2015; Vasquez-Limeta & Loncarek, 2021). This allows centrosomes to organise more MTs to promote efficient mitotic spindle assembly. The mitotic protein kinase Polo/PLK1 plays a particularly important part in centrosome maturation (Lane & Nigg, 1996; Dobbelaere *et al*, 2008; Haren *et al*, 2009; Lee & Rhee, 2011; Conduit *et al*, 2014a; Woodruff *et al*, 2015b; Ohta *et al*, 2021), and the conserved Spd-2/CEP192 family of proteins help recruit Polo/PLK1 to mitotic centrosomes (Joukov *et al*, 2014; Meng *et al*, 2015; Decker *et al*, 2011; Alvarez Rodrigo *et al*, 2019; Ohta *et al*, 2021; Wong *et al*, 2021). In flies and worms, the centrosomal Polo/PLK-1 recruited by Spd-2/SPD-2 phosphorylates the large coiled-coil proteins Cnn (flies) or SPD-5 (worms), which can then assemble into macromolecular “scaffold” structures (Conduit *et al*, 2014a; Woodruff *et al*, 2015a; Feng *et al*, 2017; Woodruff *et al*, 2017; Cabral *et al*, 2019; Ohta *et al*, 2021). These scaffolds appear to give mechanical strength to the mitotic PCM (Lucas & Raff, 2007; Mittasch *et al*, 2020) and also help to recruit many other PCM “client” proteins to the assembling mitotic centrosome (Woodruff *et al*, 2014; Raff, 2019).

In typical somatic cells the two mitotic centrosomes need to grow to approximately the same size, as mitotic centrosome size asymmetry can lead to asymmetric spindle assembly and so to defective chromosome segregation (Gasic *et al*, 2015; Meraldi, 2016). How centrosome growth is regulated in somatic cells is unclear, but in early *C. elegans* embryos, mitotic centrosome size appears to be set by a limiting pool of the PCM-scaffolding protein SPD-2 (Decker *et al*, 2011; Zwicker *et al*, 2014). The total embryonic pool of SPD-2 is thought to remain constant throughout the early cell division cycles, so the centrosomes halve in size after each round of cell division as the limiting pool of SPD-2 is divided equally amongst the exponentially increasing number of centrosomes. This concept of a “limiting pool” of an organelle building block is potentially a powerful mechanism with which to regulate organelle size, as it allows size to be set without the need for a specific size-measuring mechanism (Goehring & Hyman, 2012). It is unclear, however, if such a mechanism sets centrosome size in other cell types.

We recently developed methods to measure the growth kinetics of the Spd-2/Polo/Cnn mitotic PCM scaffold in living *Drosophila* syncytial blastoderm embryos—where we can simultaneously track tens-to-hundreds of mitotic centrosomes as they rapidly and near-synchronously assemble during S-phase in preparation for mitosis (which, in these embryos, directly follows S-phase without any Gap phase) (Foe & Alberts, 1983). These studies revealed that mitotic PCM scaffold growth was surprisingly complicated. The centrosomal levels of Polo, Spd-2 and Cnn all started to increase at the start of S-phase, but whereas Cnn levels continued to rise and/or plateau as the embryos entered mitosis, the centrosomal levels of Polo and Spd-2 started to decrease before the entry into mitosis (Wong *et al*, 2021) (Figure 1A,B). Thus, the components of the mitotic PCM scaffold exhibit different growth kinetics, making it hard to use these proteins to define centrosome “size” at any particular point in the cell cycle. We wondered whether the growth kinetics of the PCM-client proteins might give some insight into centrosome size regulation in these embryos.

**Figure 1.**
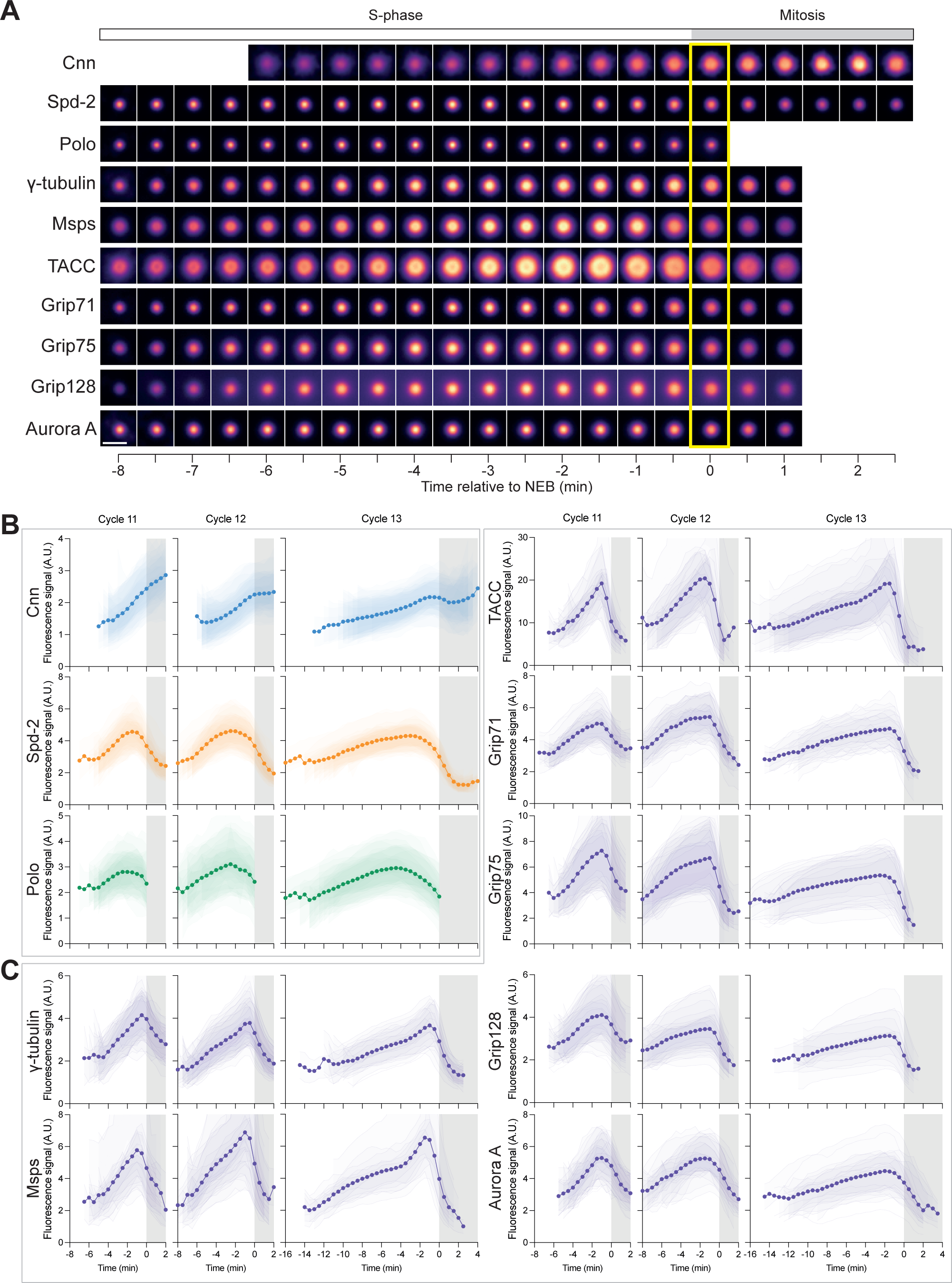
Analysis of centrosome growth kinetics. **(A)** Images show how the centrosomal fluorescence intensity of several different centrosome proteins vary during NC12 in a representative embryo (data time-normalised to NEB at t=0, *yellow box*). The images were obtained by averaging the fluorescence intensity distribution of all of the centrosomes in a single embryo at each timepoint. Note that for technical reasons not all of the centrosome proteins could be followed for the full time period. **(B,C)** Graphs show how the mean centrosomal fluorescence intensity (±SD of the data in each individual embryo shown in reduced opacity) of the PCM-scaffolding proteins Cnn, Spd-2 and Polo (B) and several PCM-client proteins (C) changes over time during NC11, 12, and 13. All individual embryo tracks were aligned to NEB (t=0). The white parts of the graphs represent S-phase, and the grey parts represent mitosis. N=7-15 embryos analysed at each nuclear cycle for each marker with a total of n=∼200-400, ∼400-800, or ∼600-1200 total centrosomes analysed at NC11, 12 and 13, respectively. Note that the data for the PCM scaffold proteins (B) was shown previously (Wong *et al*, 2021), but is reproduced here to allow comparison to the PCM client proteins (C).

Here we examine the recruitment kinetics of 7 PCM-client proteins. We find that they all accumulate at centrosomes during S-phase, reach their maximal levels just prior to mitosis, and then their centrosomal levels start to decline. Unlike the situation in early worm embryos, the mitotic centrosomes in these fly embryos grow to a relatively consistent maximal size during nuclear cycles (NC) 11, 12 and 13. This size homeostasis seems to be enforced by an inverse relationship between the centrosome growth rate and growth period. In NC11, S-phase is short and the centrosomes grow quickly for a short time; in NC13 S-phase is longer and the centrosomes grow more slowly but for a longer time. In this way, the centrosomes grow to a similar size at each cycle, irrespective of cycle length or the number of centrosomes in the embryo. The core Cdk/Cyclin cell cycle oscillator appears to play an important part in setting centrosome size in these embryos, as it can reciprocally influence the rate and period of mitotic centrosome growth.

## Results

### The dynamics of mitotic centrosome growth in the early *Drosophila* embryo

The mitotic centrosomes in early *Drosophila* embryos start to grow (i.e. accumulate PCM) during S-phase, as the embryos prepare for the next round of mitosis. We previously showed that the recruitment dynamics of the key mitotic PCM scaffolding proteins Spd-2, Polo and Cnn during NC11-13 were surprisingly complicated (Wong *et al*, 2021) (Figure 1A,B). To examine how PCM-client proteins were recruited to growing mitotic centrosomes we quantified the centrosomal fluorescence levels of γ-tubulin-GFP, Msps-GFP, GFP-TACC, Grip71-GFP, Grip75-GFP, Grip128-GFP, and Aurora A-GFP in living embryos as they proceeded through NC11-13 (Figure 1A,C). In all the experiments we report here, we therefore use the amount of a protein recruited, measured by fluorescence intensity, to define centrosome size. We obtained similar results, however, if we used centrosome area (measured from 2D projections of Z-stacks through the entire centrosome volume) as a measure of centrosome size (Figure S1).

The centrosomal levels of all seven PCM-client proteins increased during S-phase but, to our surprise, their levels started to rapidly decline shortly before the embryos entered mitosis (*grey* area on the graphs in Figure 1B,C). Although this behaviour was somewhat reminiscent of the behaviour of the Polo and Spd-2 scaffold proteins, the shape of the PCM client growth curves were quite distinct to those of Polo and Spd-2, suggesting that client protein recruitment dynamics do not simply follow Polo/Spd-2 recruitment dynamics. Moreover, the centrosomal levels of the PCM-clients appeared to decline only after the levels of Polo and Spd-2 had already started to decline (Figure S2A); this relative timing was confirmed in embryos co-expressing Spd-2-mCherry with either γ-tubulin-GFP or GFP-TACC (Figure S2B). Thus, all these PCM-client proteins exhibit recruitment dynamics that are distinct from any of the PCM scaffolding proteins, and they are all maximally recruited to centrosomes just prior to NEB, with their centrosomal levels already decreasing as the embryos enter mitosis proper.

The PCM-client recruitment profiles broadly fell into two classes that were most clearly defined by their distinct behaviours during NC13. The centrosomal levels of Grip71, Grip75, Grip128, and Aurora A tended to increase steadily through most of NC13, whereas TACC, Msps and γ-tubulin exhibited a noticeable increase in their recruitment rate towards the end of S-phase, shortly before their recruitment levels peaked (compare NC13 graphs in Figure 1B). This difference was also obvious if we used centrosome area as a measure of centrosome size (Figure S1). We conclude that PCM client proteins can be recruited to centrosomes in at least two different ways.

### Centrosome growth rate and growth period are inversely correlated so that centrosomes grow to a similar size at NC11-13

To quantify and compare the growth parameters of all the PCM-scaffold and PCM-client proteins we calculated each protein’s average initial and final centrosome-fluorescence intensity and their average rate and period of growth at each division cycle (Figure 2). Strikingly, centrosomes grew to approximately the same maximum size at each successive cycle (“peak intensity” graphs in *green* boxes, Figure 2), and this was also observed when we used centrosome area as a measure of size (“Peak area” graphs in *green* boxes, Figure S3). For some of these proteins there was a slight downward trend in their maximal fluorescence intensity across successive cycles, but this was often not statistically significant, was relatively modest—being nowhere near as large as the ∼50% decrease observed after each cell division in early *C. elegans* embryos (Decker *et al*, 2011)—and this decrease was often not observed for these proteins if we used centrosome area as a measure of centrosome size (Figure S3). We conclude that mitotic centrosomes grow to approximately the same maximal size during NC11-13. Interestingly, metaphase spindle size, measured either by the sum of the fluorescence-intensity of the spindle microtubules or by spindle length, decreased modestly, but significantly, at each successive cycle, indicating that centrosome size and metaphase spindle size do not scale proportionally in these embryos (Figure S4).

**Figure 2.**
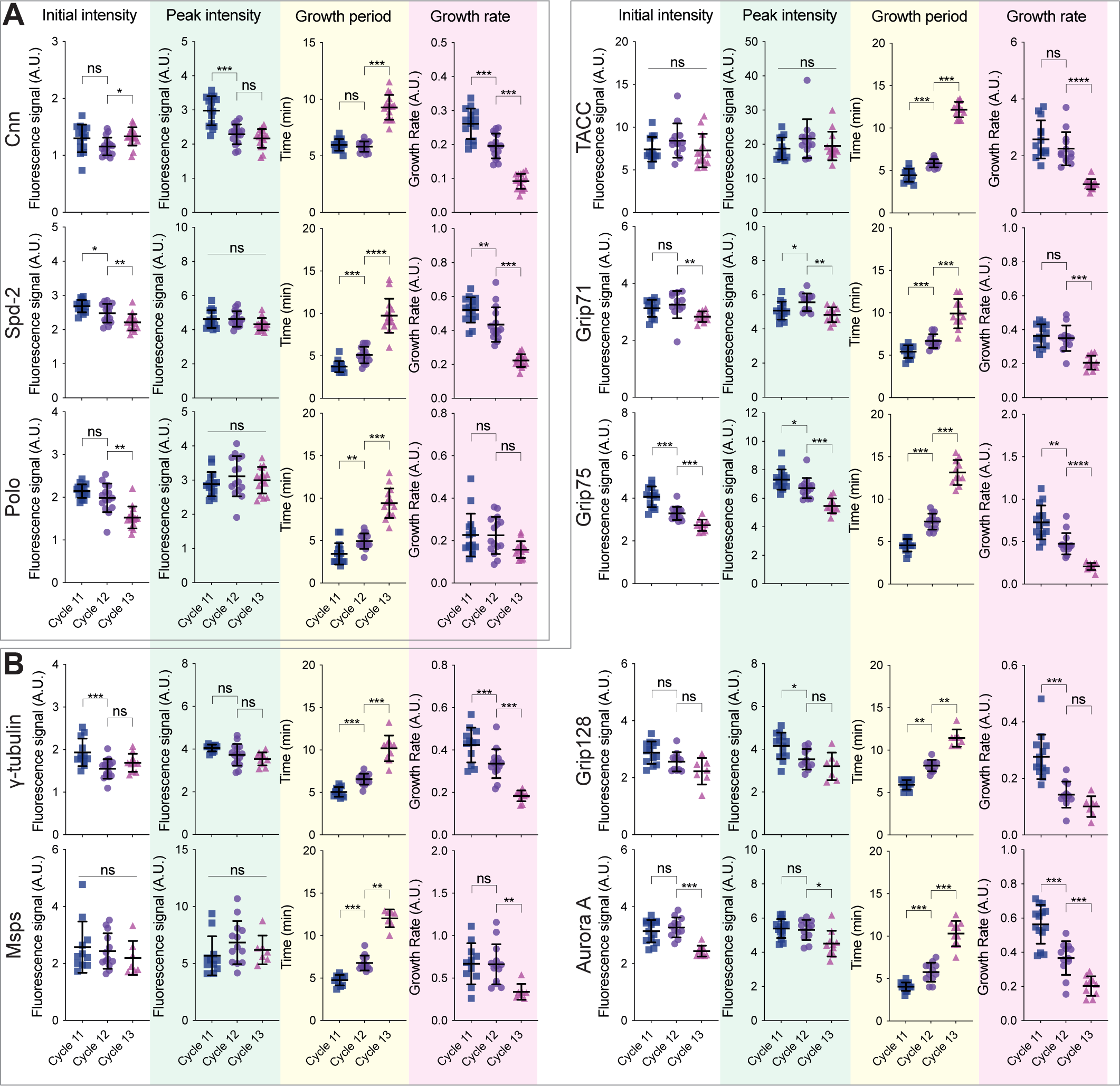
Analysis of centrosome growth parameters during NC11, 12 and 13. Scatter plots show the mean (±SD) initial fluorescent intensity (left graphs for each protein), peak fluorescent intensity (boxed in *green*), growth period (boxed in *yellow*), and growth rate (boxed in *pink*) of centrosomes during NC11, 12, or 13 for the PCM-scaffolding proteins **(A)** and the PCM-client proteins **(B)**. Each data point represents the average of all the centrosomes in an individual embryo (calculated from the data shown in Figure 1). Statistical comparisons used either an ordinary one-way ANOVA (Gaussian-distributed and variance-equal), a one-way Welch ANOVA (Gaussian-distributed and variance-unequal), or a Kruskal-Wallis’s test (non-Gaussian-distributed). If significant, multiple testing was performed using either Tukey-Kramer’s test (Gaussian-distributed and variance-equal), Games-Howell’s test (Gaussian-distributed and variance-unequal), or Mann-Whitney’s U test (non-Gaussian-distributed) (*: P<0.05, **: P<0.01, ***: P<0.001, ****: P<0.0001, ns: not significant). Gaussian distribution was tested using D’Agnostino and Pearson’s test. Variance homogeneity was tested using Levene W test.

Although the PCM-client proteins grew to a relatively consistent size at NC11-13, their growth rate and growth period varied significantly at each NC. In general, as S-phase length increased, the centrosome growth period tended to increase (graphs in *yellow* boxes, Figure 2), but the centrosome growth rate tended to decrease (graphs in *pink* boxes, Figure 2). This was also true if we used centrosome area to measure the growth rate and period, rather than fluorescence intensity (Figure S3). Thus, centrosome size homeostasis in these embryos appears to be enforced by an inverse-linear relationship between the centrosome growth rate and growth period (*left graphs* for each protein, Figure 3).

**Figure 3.**
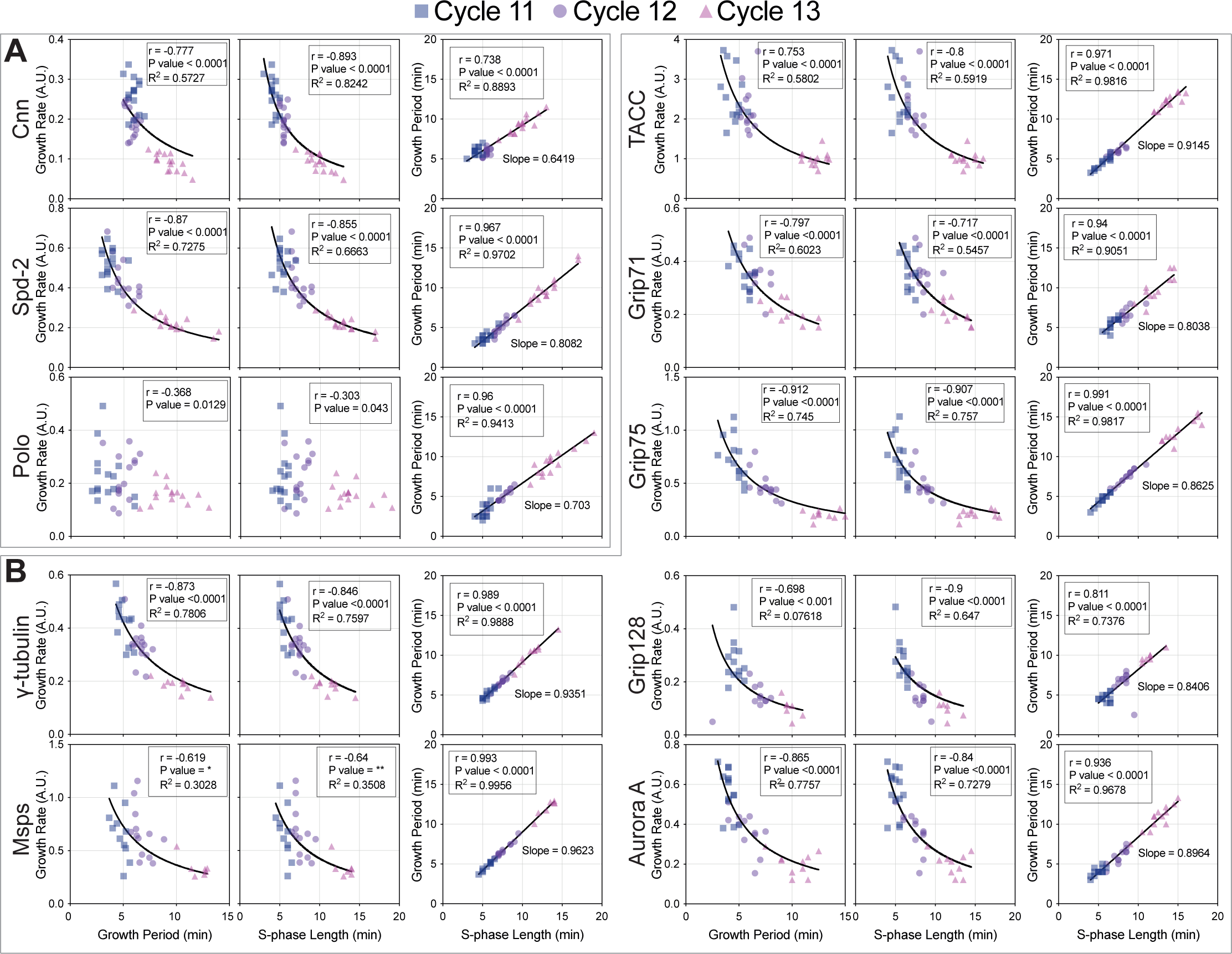
Analysis of the strength of correlation between various centrosome growth parameters during NC11, 12 and 13. Scatter plots show the correlation between the centrosome growth rate and period (left graphs for each protein), growth rate and S-phase length (middle graphs for each protein), and growth period and S-phase length (right graphs for each protein) for the centrosome scaffold **(A)** and centrosome client **(B)** proteins. Each data point represents an individual embryo at either NC11 (*deep purple squares*), NC12 (*light purple dots*) and NC13 (*pink triangles*) (calculated from the data shown in Figure 1). Lines indicate mathematically regressed best fits for inverse-linear (left and middle graphs) and linear (right graphs) correlations. The goodness of fit (R^2^), strength of correlation (r) and the statistical significance (P-value) are indicated and were calculated in custom Python scripts and GraphPad Prism by either Pearson test (bivariate Gaussian-distributed) or Spearman test (bivariate non-Gaussian distributed data). Bivariate Gaussian distribution was tested by Henze-Zirkler test. Note that for Polo, the correlation between the centrosome growth rate and either the centrosome growth period or S-phase length did not fit an inverse-linear function well, although the trend was still significant (p<0.05). This suggests that this relationship may be more complicated than for the other proteins, perhaps because Cdk/Cyclins and Polo influence each other’s behaviour in multiple ways.

### The rate of Cdk/Cyclin activation influences centrosome growth rate and period

The rapid nuclear divisions in these early embryos are driven by the activity of the core Cdk/Cyclin cell cycle oscillator (CCO). The rate at which Cdk/Cyclins are activated during each NC gradually slows during NC11-13 leading to the gradual extension of S-phase length (Farrell & O’Farrell, 2014). We speculated, therefore, that the gradual slowing of Cdk/Cyclin activation might be responsible for decreasing the centrosome growth rate and increasing the centrosome growth period at successive cycles. As the rate of Cdk/Cyclin activation is a major determinant of S-phase length in these embryos (Farrell & O’Farrell, 2014), we examined how the centrosome growth rate and growth period correlated with S-phase length. For all of the PCM-clients examined here there was a clear inverse-linear correlation between S-phase length and the centrosome growth rate (*middle graphs* for each protein, Figure 3) and a strong linear correlation between S-phase length and the centrosome growth period (*right graphs* for each protein, Figure 3).

To directly test whether the rate of Cdk/Cyclin activation influenced the PCM growth rate and growth period, we examined the recruitment dynamics of the PCM-clients γ-tubulin and TACC in embryos in which we halved the genetic dose of *Cyclin B* (hereafter *CycB^1/2^* embryos). This slows Cdk/Cyclin activation and so extends S-phase length (Aydogan *et al*, 2022) (Figure 4A-D). In *CycB^1/2^* embryos the growth period of both PCM proteins was increased, but their growth rate was slowed, so that the centrosomes grew to nearly the same size as controls (Figure 4A-D). This illustrates the homeostatic nature of centrosome growth in these embryos: the reduction in Cyclin B levels leads to a decrease in the centrosome growth rate, but also to an increase in the centrosome growth period, so that centrosomes grow to a near-normal size even though cell cycle length is altered. Thus, Cdk/Cyclin activity influences both the centrosome growth rate and growth period.

**Figure 4.**
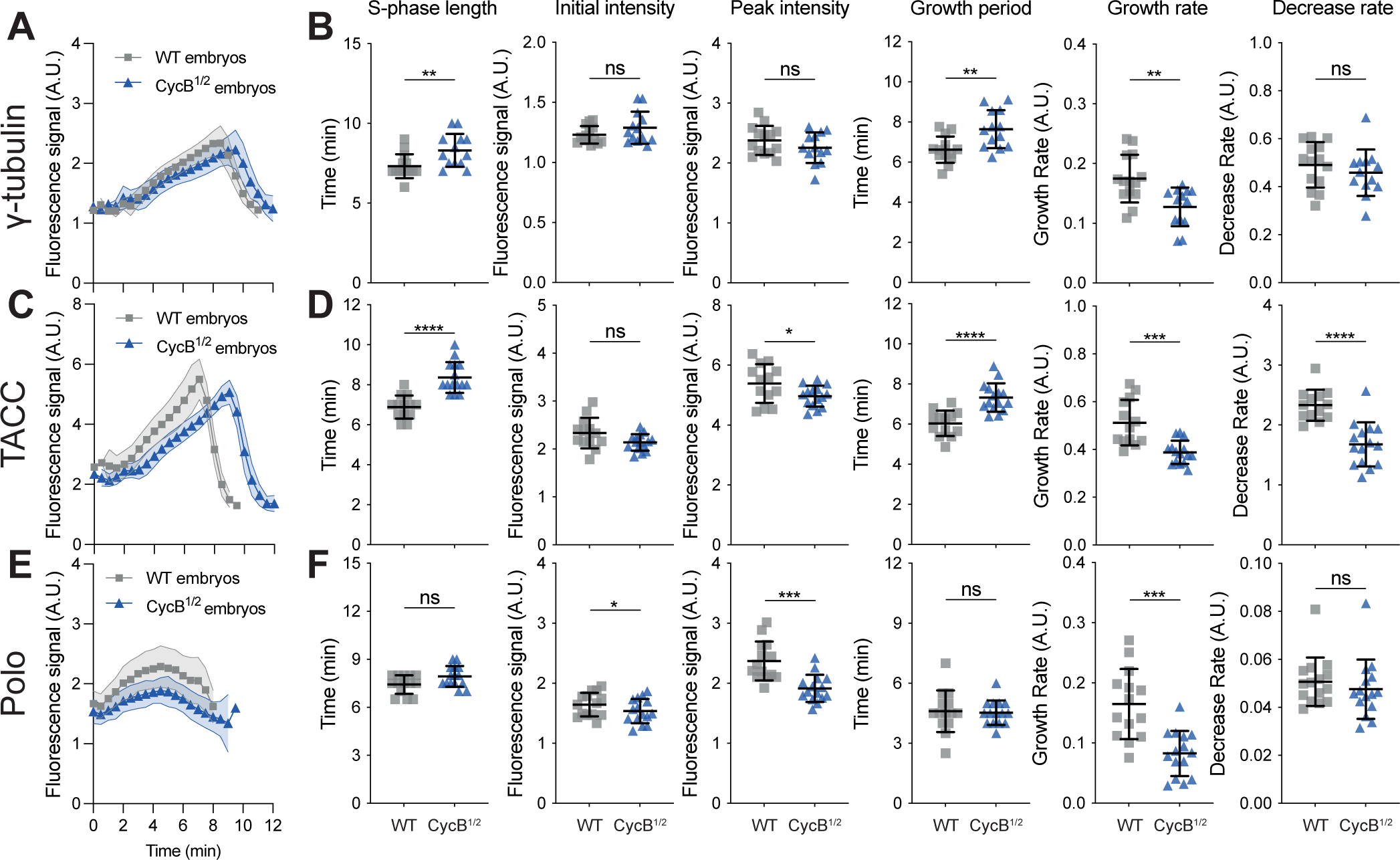
Analysis of centrosomal growth kinetics in *CycB^1/2^* embryos. **(A,C,E)** Graphs show the mean (±SD) centrosomal growth dynamics of γ-tubulin-GFP (A), GFP-TACC (C), and Polo-GFP (E) in embryos laid by either control wildtype females (*grey* lines) or heterozygous *CyclinB^+/-^*females (*CycB^1/2^* embryos) (*blue* lines). As S-phase length was usually extended in *CycB^1/2^* embryos, these data were aligned to centrosome separation (t=0), which occurs at the start of S-phase, rather than to NEB. N=10-15 embryos and a total of n=∼500-800 centrosomes were analysed for each condition. **(B,D,F)** Scatter plots compare various cell cycle and centrosome growth parameters in either WT (*grey*) or *CycB^1/2^* embryos (*blue*). Statistical significance was assessed using an unpaired t-test (*: P<0.05, **: P<0.01, ***: P<0.001, ****: P<0.0001, ns: not significant).

### Cdk/Cyclins appear to phosphorylate Spd-2 to reduce its centrosomal recruitment and/or maintenance during mitosis

How might Cdk/Cyclin activity influence the PCM growth period? Perhaps the simplest hypothesis is that Cdk/Cyclins can directly phosphorylate one or more of the PCM scaffold proteins and/or client proteins to decrease their affinity for each other as the embryos prepare to enter mitosis. In such a scenario, a threshold level of Cdk/Cyclin activity could effectively switch off mitotic centrosome growth, so centrosomes would grow for a longer period in embryos where Cdk/Cyclin activation is slowed (e.g. during later NCs or when we reduce the dosage of Cyclin B).

We first wanted to test whether direct phosphorylation by Cdk/Cyclins might influence the behaviour of the PCM scaffold. We mutated potential Cdk/Cyclin phosphorylation sites (Ser/Thr-Pro motifs mutated to Ala-Pro) in the PCM scaffold proteins Spd-2 (20 in total) and Cnn (6 in total) (Figure S5), to generate mutant proteins (Spd-2-Cdk20A and Cnn-Cdk6A, respectively). We generated transgenic lines expressing either untagged or mNeonGreen (NG)-tagged versions of the mutant proteins, and found that all of these lines were dominant male sterile. Our preliminary analyses indicate that the expression of these proteins led to an accumulation of cytoplasmic aggregates of the mutant proteins in spermatocytes during meiosis (Figure S6) and a subsequent failure in cytokinesis—a phenomenon that will be investigated in more detail elsewhere. This dominant male sterility meant that we could only examine the behaviour of the mutant proteins in embryos laid by females carrying one copy of the transgene and two WT (untagged) copies of the endogenous gene (i.e. in the presence of significant amounts of WT, untagged protein). As controls, we therefore examined the behaviour of WT Spd-2-NG and WT NG-Cnn in embryos expressing one copy of the NG-fusion in the presence of two copies of the WT untagged endogenous gene (Figure 5A,F). We compared the recruitment kinetics of these fusion-proteins during NC12.

**Figure 5.**
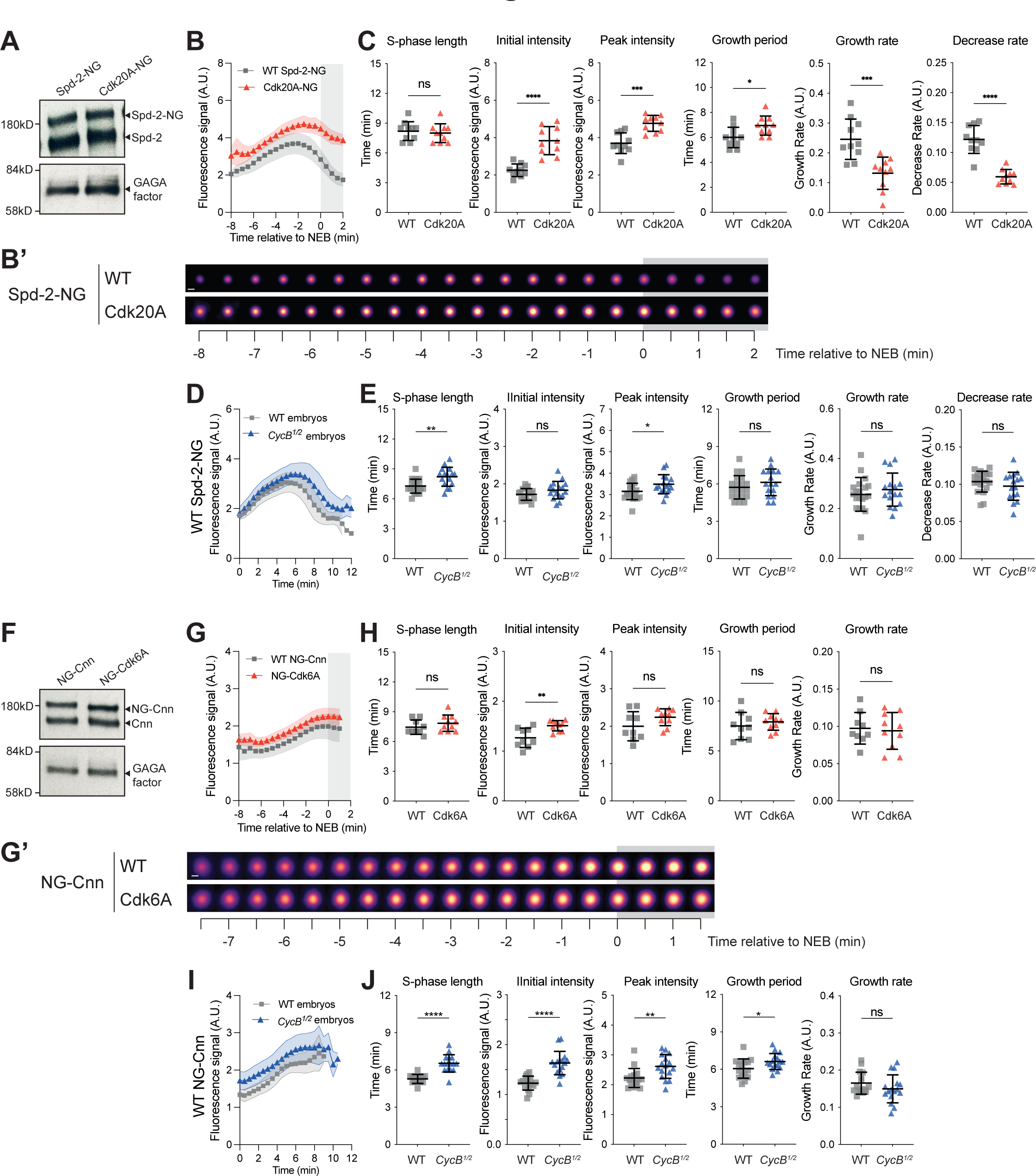
Analysis of Spd-2 and Cnn centrosome growth kinetics when their potential phosphorylation by Cdk/Cyclins is perturbed. **(A)** Western blot shows protein-levels of the endogenous untagged Spd-2 and transgenically expressed WT Spd-2-NG or the Spd-2-Cdk20A mutant that should not be phosphorylated efficiently by Cdk/Cyclins. GAGA transcription factor is shown as a loading control. **(B,B’)** Graph (B) and images of centrosomes from a representative embryo (B’) compare how the centrosomal fluorescence intensity (mean±SD in the graph) of WT Spd-2-NG or Spd-2-Cdk20A change over time during NC12. Timepoints shaded in *grey* represent mitosis. The images were obtained by averaging the fluorescence intensity of all of the centrosomes in a single embryo at each timepoint. **(C)** Scatter plots show the mean (±SD) of various cell cycle and centrosome-growth parameters derived from the data shown in (B). **(D)** Graph compares how the centrosomal fluorescence intensity (mean±SD) of WT Spd-2-NG changes over time in either WT or *CycB^1/2^* embryos. **(E)** Scatter plots show the mean (±SD) of various cell cycle and centrosome-growth parameters derived from the data shown in (D). **(F-J)** Panels show the same analyses as described in (A-E) but comparing the behaviour of WT NG-Cnn to the NG-Cnn-Cdk6A mutant. N=9-17 embryos and a total of n=∼400-1000 centrosomes were analysed for each condition. Statistical comparisons used either an ordinary one-way ANOVA (Gaussian-distributed and variance-equal), a one-way Welch ANOVA (Gaussian-distributed and variance-unequal), or a Kruskal-Wallis’s test (non-Gaussian-distributed). If significant, multiple testing was performed using either Tukey-Kramer’s test (Gaussian-distributed and variance-equal), Games-Howell’s test (Gaussian-distributed and variance-unequal), or Mann-Whitney’s U test (non-Gaussian-distributed) (*: P<0.05, **: P<0.01, ***: P<0.001, ****: P<0.0001, ns: not significant). Gaussian distribution was tested using D’Agnostino and Pearson’s test. Variance homogeneity was tested using Levene W test.

Although WT Spd-2-NG and Spd-2-Cdk20A-NG were present in embryos at similar levels (Figure 5A), the centrosomal levels of the mutant protein were elevated, its growth period was extended, while the rate at which its centrosomal levels decreased as the embryos entered mitosis was dramatically slowed (Figure 5B,B’,C). This behaviour is consistent with the possibility that Cdk/Cyclins normally phosphorylate Spd-2 to decrease the efficiency of its centrosomal recruitment and/or maintenance in the run-up to mitosis. Interestingly, such a mechanism could also help to explain why centrosomal levels of WT Spd-2 normally decrease as the embryo prepares to enter mitosis (Figure 1B). In support of this possibility, WT Spd-2-NG was recruited more strongly to centrosomes in *CycB^1/2^*embryos (Figure 5D,E). This effect was not as striking as that seen with Spd-2-Cdk20A, presumably because reducing the dosage of Cyclin B only reduces WT Spd-2 phosphorylation by Cdk/Cyclins, whereas this phosphorylation is potentially abolished by the Spd-2-Cdk20A mutations. Together, these data suggest that Cdk/Cyclins directly phosphorylate Spd-2 in the run-up to mitosis to help reduce Spd-2’s centrosomal recruitment and/or maintenance.

WT NG-Cnn and NG-Cnn-Cdk6A were present in embryos at similar levels (Figure 5F), and the centrosomal levels of the mutant protein increased slightly compared to WT NG-Cnn (Figure 5G). This difference was not as large as that seen for the Spd-2-Cdk20A-NG mutant, and most growth parameters of NG-Cnn-Cdk6A were not statistically different to WT (Figure 5G,H). We also observed a modest increase in the centrosomal levels of WT NG-Cnn in *CycB^1/2^* embryos (Figure 5I,J), but note that this could be explained, at least in part, by the increase in centrosomal Spd-2 levels in *CycB^1/2^* embryos (Figure 5D,E), as Spd-2 helps recruit Cnn to centrosomes (Conduit *et al*, 2014b). Thus, any potential direct phosphorylation of Cnn by Cdk/Cyclins does not appear to strongly influence centrosomal Cnn recruitment and/or maintenance.

### Cdk/Cyclins appear to phosphorylate Spd-2 and Cnn to reduce their ability to recruit and/or maintain γ-tubulin at centrosomes

We next wanted to examine whether preventing the Cdk/Cyclin-dependent phosphorylation of Spd-2 and/or Cnn could influence the centrosomal recruitment of any PCM clients. We focused on γ-tubulin because both the Spd-2/CEP192 and Cnn/CDK5RAP2 family of proteins have been implicated in recruiting γ-tubulin to centrosomes (Gomez-Ferreria *et al*, 2007; Zhu *et al*, 2008; O’Rourke *et al*, 2014; Fong *et al*, 2008; Choi *et al*, 2010; Tovey *et al*, 2021; Ohta *et al*, 2021). In addition, we found that reducing the levels of Spd-2 or Cnn in these early embryos reduced the centrosomal recruitment of γ-tubulin in distinct ways, suggesting that both proteins contribute independently to this process (Figure 6). We expressed a γ-tubulin-GFP fusion protein in embryos co-expressing untagged versions of either WT-Spd-2 or Spd-2-Cdk20A or WT-Cnn or Cnn-Cdk6A. The expression of the mutant proteins did not appear to dramatically perturb the centrosomal recruitment of γ-tubulin-GFP, except that the rate at which γ-tubulin-GFP left the centrosome as the embryos entered mitosis was reduced in both mutants compared to WT (Figure 7). This phenotype was subtle, but it was statistically significant, and it seems likely that the presence of large amounts of WT Spd-2 and Cnn in the mutant embryos (Figure 5A,F) would help to mask the potential severity of this phenotype. These observations suggest that Cdk/Cyclins can normally phosphorylate Spd-2 and Cnn to decrease their ability to recruit and/or maintain centrosomal γ-tubulin as the embryos prepare to enter mitosis.

**Figure 6.**
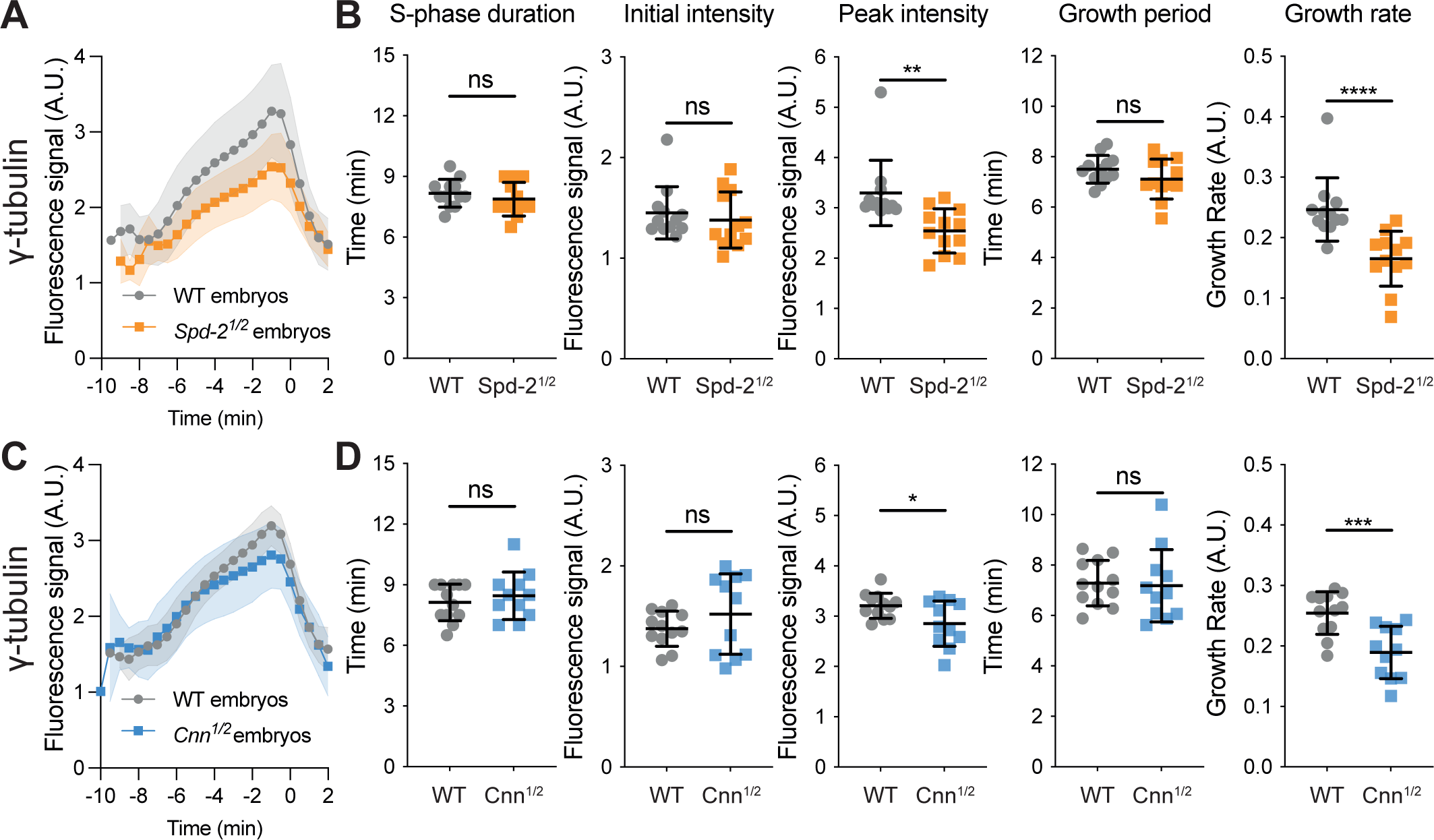
Analysis of γ-tubulin-GFP centrosome growth kinetics in embryos with reduced levels of Spd-2 or Cnn. **(A,C)** Graphs compare how the mean (±SD) centrosomal fluorescence intensity of γ-tubulin-GFP changes over time during NC12 in embryos laid by either WT mothers or mothers heterozygous for *Spd-2* or *cnn* mutations (*Spd-2^1/2^* or *Cnn^1/2^* embryos, respectively). Individual embryo tracks were aligned to NEB (t=0). N=10-15 embryos and a total of n=∼500-800 centrosomes were analysed for each condition. **(B,D)** Scatter plots compare the mean (±SD) of various cell cycle and centrosome-growth parameters in WT and *Spd-2^1/2^* or *Cnn^1/2^* embryos. Statistical comparisons used either an ordinary one-way ANOVA (Gaussian-distributed and variance-equal), a one-way Welch ANOVA (Gaussian-distributed and variance-unequal), or a Kruskal-Wallis’s test (non-Gaussian-distributed). If significant, multiple testing was performed using either Tukey-Kramer’s test (Gaussian-distributed and variance-equal), Games-Howell’s test (Gaussian-distributed and variance-unequal), or Mann-Whitney’s U test (non-Gaussian-distributed) (*: P<0.05, **: P<0.01, ***: P<0.001, ****: P<0.0001, ns: not significant).. Gaussian distribution was tested using D’Agnostino and Pearson’s test. Variance homogeneity was tested using Levene W test.

**Figure 7.**
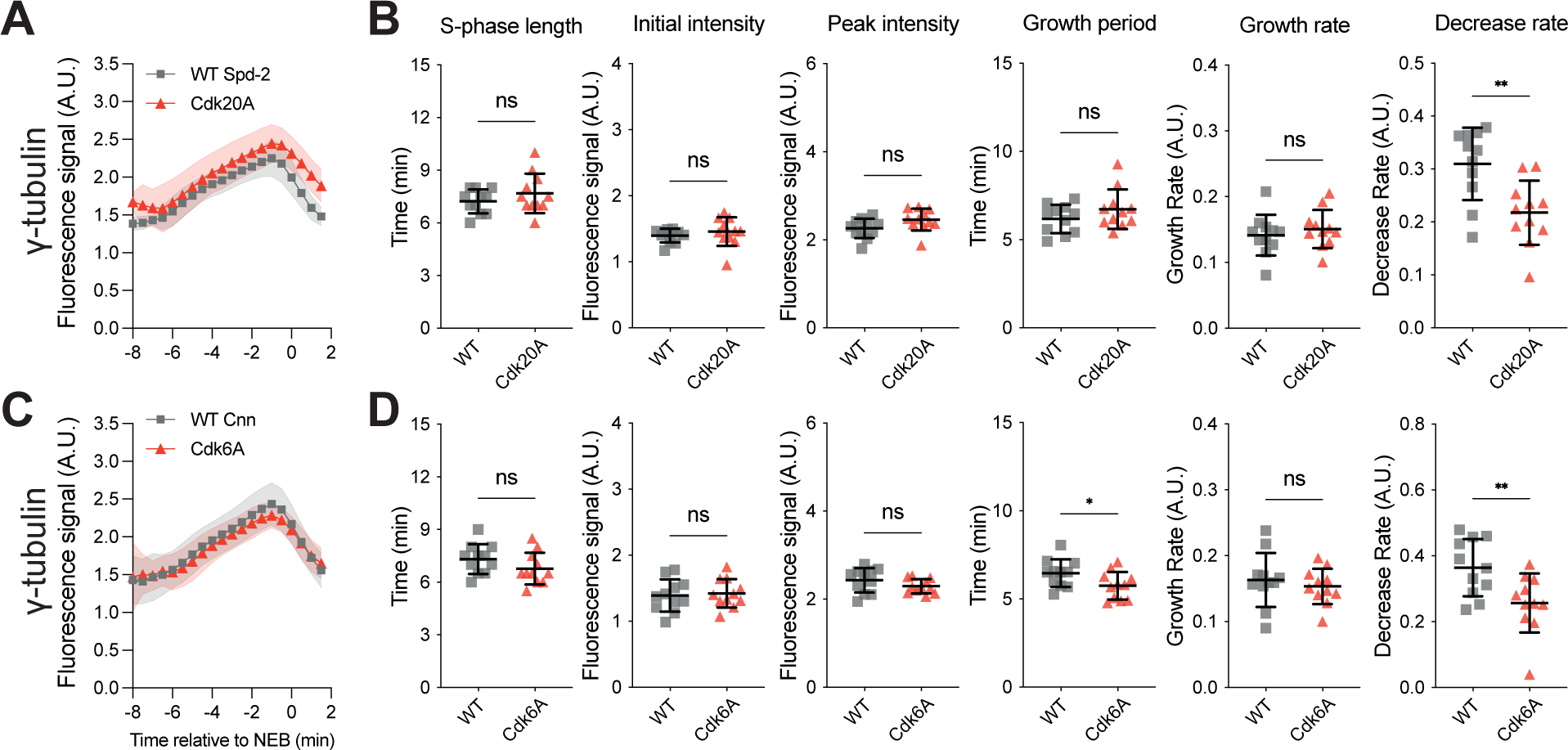
Analysis of γ-tubulin-GFP centrosome growth kinetics in embryos expressing mutant forms of Spd-2 or Cnn that should not be phosphorylated by Cdk/Cyclins efficiently. **(A,C)** Graphs compare how the mean centrosomal fluorescence intensity (±SD) of γ-tubulin-GFP changes over time during NC12 in WT embryos expressing untagged versions of either WT or mutant forms of Spd-2 (Spd-2-Cdk20A) (A) or Cnn (Cnn-Cdk6A) (C) that should not be phosphorylated by Cdk/Cyclins efficiently. Individual embryo tracks were aligned to NEB (t=0). 10-15 embryos and a total of 500-800 centrosomes were analysed for each condition. **(B,D)** Scatter plots compare the mean (±SD) of various cell cycle and centrosome-growth parameters in embryos expressing the WT and mutant proteins. Statistical comparisons used either an ordinary one-way ANOVA (Gaussian-distributed and variance-equal), a one-way Welch ANOVA (Gaussian-distributed and variance-unequal), or a Kruskal-Wallis’s test (non-Gaussian-distributed). If significant, multiple testing was performed using either Tukey-Kramer’s test (Gaussian-distributed and variance-equal), Games-Howell’s test (Gaussian-distributed and variance-unequal), or Mann-Whitney’s U test (non-Gaussian-distributed) (*: P<0.05, **: P<0.01, ns: not significant). Gaussian distribution was tested using D’Agnostino and Pearson’s test. Variance homogeneity was tested using Levene W test.

Finally, we wanted to examine whether mutating the potential Cdk/Cyclin phosphorylation sites in γ-tubulin (7 in total) perturbed its centrosomal recruitment dynamics. Unfortunately, a NG-fusion to this mutant protein did not detectably localise to centrosomes, so we were unable to assess whether Cdk/Cyclins directly phosphorylate γ-tubulin to reduce its affinity for the PCM scaffold.

### Cdk/Cyclin activity influences the amount of Polo recruited to centrosomes

Our studies so far suggest mechanisms that could help to explain how Cdk/Cyclin activity influences the centrosome growth period. But how might Cdk/Cyclin activity influence the centrosome growth rate? We noticed that for the majority of the centrosome proteins we analysed, the amount of protein initially recruited to centrosomes at the start of NC11-13 was relatively constant at successive cycles and exhibited no clear trend, and this was also true when centrosome area was used as a measure of centrosome size (left graphs, Figure 2 and Figure S3). The initial centrosomal levels of Polo-GFP, however, exhibited a downward trend across NC11-13, and this was also seen when centrosome area was used as a measure of centrosome size (left graphs, Figure 2 and Figure S3). The activation of Cdk/Cyclin kinases and Polo/PLK1 kinases through the cell cycle are intimately linked (Lindqvist *et al*, 2009; Crncec & Hochegger, 2019) and Polo/PLK1 is a major driver of mitotic centrosome growth, so we wondered whether Cdk/Cyclins might influence the rate of centrosome growth by influencing Polo recruitment to centrosomes.

To test this possibility, we examined the centrosomal recruitment dynamics of Polo-GFP in *CycB^1/2^* embryos. Strikingly, whereas the PCM clients γ-tubulin and TACC reciprocally adjusted their growth rate and growth period so that the centrosomes ultimately recruited near-normal amounts of each protein in *CycB^1/2^* embryos, this was not the case for Polo-GFP, whose centrosomal levels were strongly reduced in *CycB^1/2^* embryos (Figure 4E,F). We conclude that Cdk/Cyclin activity normally promotes Polo recruitment to and/or maintenance at centrosomes, potentially helping to explain how Cdk/Cyclin activity influences the centrosome growth rate.

## Discussion

Here we examine how centrosome size is regulated in syncytial blastoderm *Drosophila* embryos. We find that centrosomes grow to a similar size at nuclear cycles 11-13, and this size homeostasis is enforced, at least in part, by an inverse-linear relationship between the centrosome growth rate and growth period: centrosomes grow rapidly, but for a short period, in cycle 11, and then more slowly, but for a longer period, at later cycles. As a result, the centrosomes grow to a relatively consistent size no matter the length of the nuclear cycle or the number of centrosomes present in the embryo. These observations suggest that, in contrast to early worm embryos (Decker *et al*, 2011; Goehring & Hyman, 2012), centrosome size is not set by the cytoplasmic depletion of a limiting component in syncytial fly embryos. Given that fly and worm embryos use a similar set of proteins to build their mitotic centrosomes (Conduit *et al*, 2015; Pintard & Bowerman, 2019), it is perhaps surprising that the mechanisms regulating centrosome size appear to be so different. Clearly more work is required to resolve this puzzling paradox.

The inverse-linear relationship we observe between the centrosome growth rate and growth period in fly embryos seems to be enforced, at least in part, by the Cdk/Cyclin cell cycle oscillator (CCO). In early *Drosophila* embryos, the rate of CCO activation at the start of each nuclear cycle gradually slows during cycles 10-14 (Farrell & O’Farrell, 2014). This leads to an increase in S-phase length (Foe & Alberts, 1983; Farrell & O’Farrell, 2014) and, as we show here, to an increase in the centrosome growth period but a decrease in the centrosome growth rate. Our findings suggest that the CCO influences the parameters of centrosome growth in multiple ways (summarised in Figure 8).

**Figure 8.**
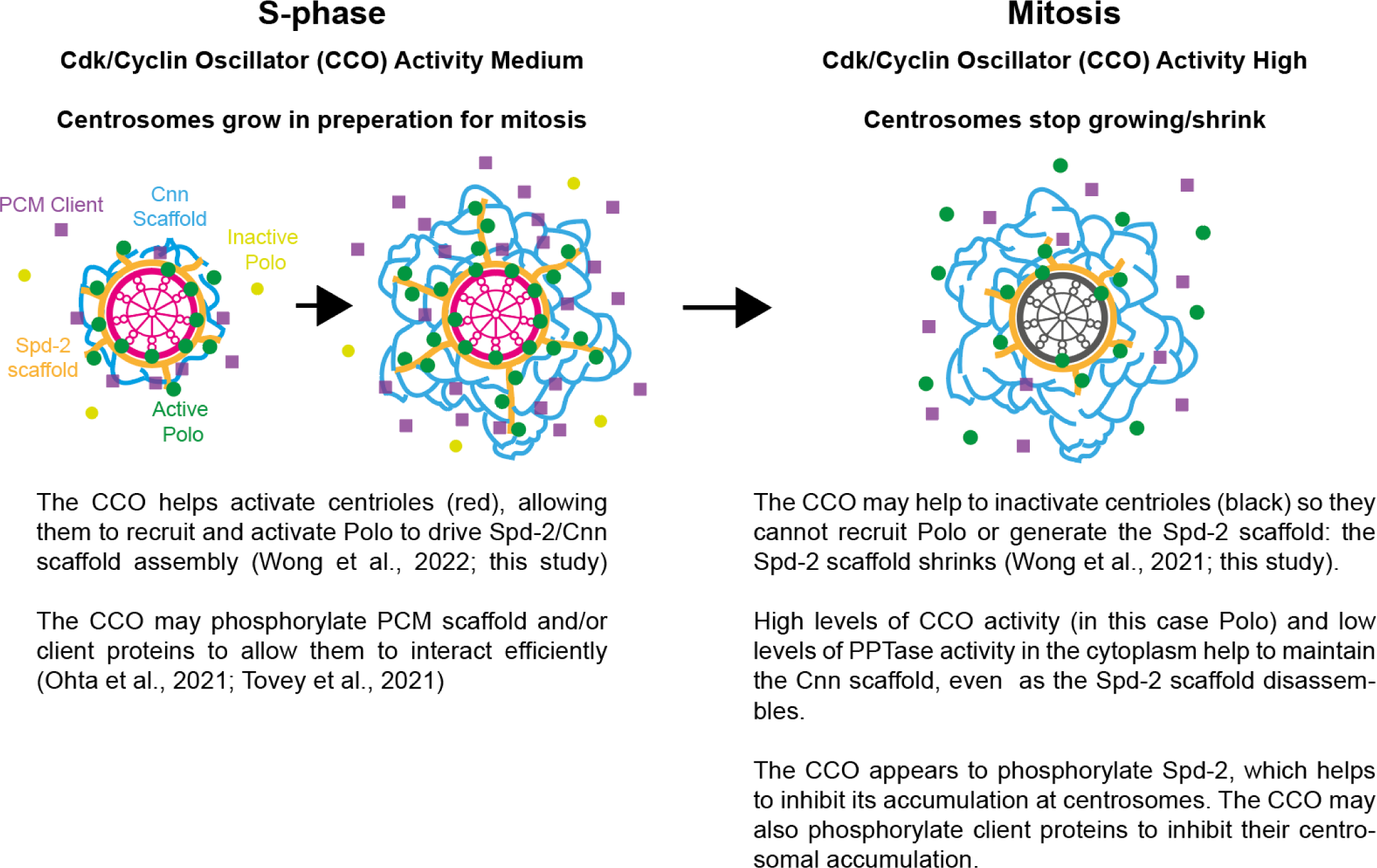
Schematic summary of how Cdk/Cyclin oscillator (CCO) activity appears to regulate mitotic centrosome growth in *Drosophila* embryos. In *syncytial* Drosophila embryos, Cdk/Cyclin activity is high enough at the start of S-phase that DNA synthesis is initiated at all replication origins, which allows the DNA to replicate quickly (Farrell & O’Farrell, 2014). This level of Cdk/Cyclin activity (which in more typical somatic cells would only be present towards the end of S-phase/early G2) is sufficient to activate the centrioles to allow them to recruit Polo and initiate Spd-2/Cnn scaffold assembly. Mitotic centrosomes start to grow, driven by the expansion of the scaffold, but also potentially by the phosphorylation of scaffold and/or client proteins to increase their interactions (Ohta *et al*, 2021; Tovey *et al*, 2021). As Cdk/Cyclin activity approaches its maximal levels at mitotic entry, however, our data suggests that Spd-2—and possibly Cnn and some client proteins—can be further phosphorylated (either directly by Cdk/Cyclins, as we show here for Spd-2, or perhaps by other mitotic kinases such as Polo and/or Aurora A) which decreases their interactions and so helps to suppress centrosome growth. In addition, the centrioles in these embryos switch off their ability to recruit Polo and generate the Spd-2 scaffold (Wong *et al*, 2021). Together, these mechanisms help to ensure that the centrosomes stop growing, and in *Drosophila* embryos actually start to shrink (at least in terms of the Spd-2/Polo scaffold and PCM client proteins), as the embryos prepare to enter mitosis. In contrast, the Cnn scaffold is maintained at centrosomes during mitosis, probably because it must be dephosphorylated to disassemble (Conduit *et al*, 2014b; Feng *et al*, 2017); the high levels of cytoplasmic Polo activity during mitosis (and likely the low levels of relevant cytoplasmic phosphatase activity) mean that the Cnn scaffold is not dephosphorylated and disassembled until the end of mitosis.

First, the moderate levels of CCO activity present in syncytial *Drosophila* embryos at the start of S-phase appear to promote centrosome growth by driving Polo recruitment to the assembling mitotic centrosomes. Polo/PLK1 recruitment to centrosomes is mediated by its Polo-Box domain, which binds to phosphorylated S-S/T(P) motifs, and this binding is sufficient to partially activate the kinase (Lee *et al*, 1998; Song *et al*, 2000; Elia *et al*, 2003; Reynolds & Ohkura, 2003; Liu *et al*, 2004). We recently showed that in fly embryos S-S/T(P) motifs in the centriole protein Ana1/CEP295 (Saurya *et al*, 2016) and the PCM-scaffolding protein Spd-2/CEP192 (Dix & Raff, 2007; Giansanti *et al*, 2008) cooperate to recruit Polo first to centrioles (mainly dependent on Ana1/CEP295) and then into the expanding mitotic PCM (mainly dependent on Spd-2/CEP192) (Alvarez Rodrigo *et al*, 2019; Alvarez-Rodrigo *et al*, 2021; Wong *et al*, 2021). The function of Spd-2/CEP192 in recruiting Polo/PLK1 to the mitotic PCM is conserved in vertebrates and worms (Decker *et al*, 2011; Joukov *et al*, 2014; Meng *et al*, 2015). Although the precise S-S/T(P) motifs in Ana1 and Spd-2 required for Polo recruitment in flies have not been identified, several candidates are potential Cdk/Cyclin phosphorylation sites, as is the single PLK1-recruiting motif in worm SPD-2 (Decker *et al*, 2011). Thus, it seems plausible that Cdk/Cyclin activity might directly influence Polo recruitment to centrioles/centrosomes by influencing the phosphorylation of S-S/T(P) motifs in Ana1/CEP295 and/or Spd-2/CEP192.

Second, our studies indicate that, perhaps counter-intuitively, the higher levels of Cdk/Cyclin activity present in embryos that are about to enter mitosis inhibits centrosome growth. Multiple mechanisms probably contribute to this inhibition, and we identify two candidates here: (1) Cdk/Cyclins appear to directly phosphorylate Spd-2 to inhibit its centrosomal recruitment and/or maintenance; (2) Cdk/Cyclins appear to directly phosphorylate both Spd-2 and Cnn to inhibit their ability to recruit and/or maintain γ-tubulin at centrosomes. *A priori*, this latter mechanism might appear at odds with previous reports that the phosphorylation of Cnn and SPD-5 is required to allow these proteins to recruit

γ-tubulin (Ohta *et al*, 2021; Tovey *et al*, 2021). We suspect, however, that these previously reported phosphorylation events occur when CCO activity starts to rise in preparation for mitosis (so promoting centrosome growth), but that additional phosphorylation events occur as CCO activity increases, and these inhibit the recruitment and/or maintenance of γ-tubulin (and perhaps also of other PCM client proteins). It is well established that different levels of Cdk/Cyclin activity promote different phosphorylation events as cells progress through the cell cycle (Swaffer *et al*, 2016).

Thus, we envisage a scenario in which the moderate levels of CCO activity present in fly embryos at the start of S-phase—which would be equivalent to the levels of activity found in somatic cells during late S-phase/early G2 (Farrell & O’Farrell, 2014)—help to initiate mitotic centrosome growth, while the higher levels of CCO activity present at the start of mitosis help to inhibit centrosome growth (Figure 8). Such a system would help to establish an inverse-linear relationship between the centrosome growth rate and growth period, and so promote centrosome size homeostasis. In early nuclear cycles, for example, CCO activity rises quickly, so centrosomes grow quickly (as they recruit Polo quickly), but they only grow for a short time (as CCO activity quickly reaches the threshold required to switch off centrosome growth). At later nuclear cycles, when CCO activity rises more slowly (or in situations where we artificially slow CCO activation by reducing the dosage of Cyclin B) centrosomes would grow more slowly, but for a longer period.

We were surprised to find that in these rapidly cycling fly embryos the levels of most of the centrosome proteins we analysed here start to decline prior to mitotic entry. This behaviour appears to be unusual, perhaps suggesting that the inhibition of centrosome growth when Cdk/Cyclin activity is high is a specific adaptation of the rapidly cycling early fly embryo. Previous studies that have measured or inferred the kinetics of centrosome growth in vertebrate cells, however, report that centrosomes stop growing at about the time the cells enter mitosis (Piehl *et al*, 2004; Khodjakov & Rieder, 1999). Thus, in both fly embryos and vertebrate cells mitotic centrosomes stop growing at about the time of NEB (although, unlike in the fly embryo, centrosomes do not appear to decrease in size until later in mitosis in vertebrate cells). Thus, the inhibition of mitotic centrosome growth by high-levels of Cdk/Cyclin activity may be conserved. The situation in worm embryos is unclear as some studies report that centrosomes continue to grow during mitosis (Decker *et al*, 2011; Wueseke *et al*, 2016; Cabral *et al*, 2019), while others find that centrosome size plateaus as the embryos enter mitosis (Ohta *et al*, 2021). These differences may be due to the different proteins being monitored and/or the different methods used to measure centrosome size.

It is not clear why it might be beneficial for mitotic centrosomes to decrease in size prior to NEB in early fly embryos. Perhaps in these rapidly cycling embryos it is important to start disassembling the mitotic PCM early, as mitosis lasts only ∼3-4mins in these embryos (Foe & Alberts, 1983), so the mitotic centrosomes must very rapidly disassemble and then initiate a new round of assembly immediately after mitotic exit. Moreover, unlike worm embryos, fly embryos have strong non-centrosomal pathways of spindle-assembly (Hayward *et al*, 2014), so centrosome-MTs might be most important for breaking down the nuclear envelope (Katsani *et al*, 2008); after NEB, it may be useful to divert MT nucleating/organising resources away from centrosomes to the other pathways of spindle assembly (Petry, 2016; Prosser & Pelletier, 2017). This may help to explain our observation that centrosome size and spindle size do not appear to scale proportionally in early fly embryos, where as they do scale in early worm embryos (Greenan et al., 2010).

Finally, it is fascinating to note that in fly embryos an inverse relationship between the growth rate and growth period seems to help set both daughter centriole (Aydogan *et al*, 2018b, 2020a) and mitotic centrosome (this study) size. In fly embryos, both assembly pathways are initiated when centrioles generate a local pulse of kinase activity—a PLK4 pulse to initiate daughter centriole growth (Aydogan *et al*, 2018b, 2020a) and a PLK1 pulse to initiate mitotic centrosome growth (Wong *et al*, 2021) (this study). These pulses are normally entrained by the Cdk/Cyclin cell cycle oscillator and, in fly embryos, both pulses are initiated in S-phase when Cdk/Cyclin activity is moderate. In more typical somatic cells, we would envisage that S-phase levels of Cdk/Cyclin activity would stimulate the centriole-pulse of PLK4 to initiate centriole assembly, while G2 levels of activity would stimulate the centriole-pulse of PLK1 to initiate mitotic centrosome assembly. The pulsatile nature of PLK4 and PLK1 recruitment helps to limit organelle growth to ensure that centrioles and mitotic centrosomes grow to a consistent size. In addition, however, mitotic levels of Cdk/Cyclin activity also serve to inhibit organelle growth by phosphorylating key centriole/centrosome building blocks—Ana2/STIL to inhibit daughter centriole growth (Steinacker *et al*, 2022) and Spd-2/CEP192 to inhibit mitotic centrosome growth (this study).

Could similar principles regulate the growth of other organelles? This might seem far-fetched, as the mechanisms regulating centriole and mitotic centrosome growth have presumably evolved to generate organelles of a consistent size, whereas other organelles may need to be more responsive to cellular conditions to set their size. We note, however, that the biogenesis of many membrane bound organelles is thought to occur at local membrane contact sites where key activities are concentrated (Wu *et al*, 2018; Farré *et al*, 2019; Prinz *et al*, 2020). It will be interesting to determine if the levels of these key activities are locally pulsatile, if these pulses reciprocally influence the rate and period of organelle growth, and whether the cell cycle oscillator (or perhaps some other master oscillator, such as the circadian clock) regulates these pulses to influence organelle size and coordinate organelle biogenesis with other cellular events.

## Acknowledgements

We are grateful to members of the Raff Laboratory for advice, discussion and for critically reading the manuscript. The research was funded by a Wellcome Trust Senior Investigator Award (215523) to J.W. Raff (A. Wainman, S. Saurya) and a Cancer Research UK Oxford Centre Prize DPhil Studentship (C5255/A23225) and a Balliol Jason Hu Scholarship and a Clarendon Scholarship (to S.S. Wong).

## Author contributions

This study was conceptualised by S.S.W. and J.W.R. Investigation was done by S.S.W., S.S., and A.W. Data was analysed by S.S.W., S.S., A.W., and J.W.R. The project was supervised and administered by J.W.R. All authors contributed to the drafting and editing of the manuscript.

## Declaration of interests

The authors declare no competing interests.

## Rights Retention Statement

This research was funded in whole or in part by Wellcome (215523). For the purpose of Open Access, the author has applied a CC BY public copyright licence to any Author Accepted Manuscript (AAM) version arising from this submission.

## Supplementary Material

### Materials and Methods

#### Drosophila melanogaster stocks and husbandry

The *Drosophila* stocks used, generated and/or tested in this study are listed in Table 1. The precise stocks used in each experiment (and the relevant Figure) are listed in Table 2. Flies were maintained on *Drosophila* culture medium (0.68% agar, 2.5% yeast extract, 6.25% cornmeal, 3.75% molasses, 0.42% propionic acid, 0.14% tegosept, and 0.7% ethanol) in 8cm × 2.5cm plastic vials or 0.25-pint plastic bottles. For microscopy and immunoblot experiments, flies were placed in embryo collection cages on fruit juice plates (see below) with a drop of yeast paste. Fly handling were performed as previously described (Roberts, 1998).

**Table 1:** *Drosophila* stocks used in this study.

| Allele | Source |
| --- | --- |
| WT, Canton S | Bloomington Stock Centre |
| cnn<sup>f04547</sup> | Exelixis stock no. f04547, Exelixis Stock Centre (Harvard Medical School, Boston, MA). |
| cnn<sup>HK21</sup> | (Megraw *et al*, 1999; Vaizel-Ohayon & Schejter, 1999) |
| Ubq-NG-Cnn | (Wong *et al*, 2021) |
| Ubq-Spd-2-GFP | (Dix & Raff, 2007b) |
| Spd-2<sup>z35711</sup> | (Giansanti *et al*, 2008b) |
| Spd-2<sup>G20143</sup> | (Dix & Raff, 2007b) |
| Polo-TRAP-GFP | (Buszczak *et al*, 2007); appears to not be fully functional and is only viable as a heterozygote. |
| ncd-γ-tubulin-37c-GFP | (Hallen *et al*, 2008) |
| Ubq-Msps-GFP | (Lee *et al*, 2001) |
| Ubq-GFP-TACC | (Barros *et al*, 2005) |
| tacc<sup>stella592</sup> | (Lee *et al*, 2001) |
| Ubq-Grip71-GFP | (Conduit *et al*, 2014c) |
| Grip75-GFP | (Tovey *et al*, 2020); CRISPR Knock-in fast folded GFP |
| Grip128-GFP | (Tovey *et al*, 2020); CRISPR Knock-in fast folded GFP |
| Ubq-Aurora A-GFP | (Lucas & Raff, 2007b) |
| cycB<sup>2</sup> | (Jacobs *et al*, 1998) |
| Ubq-Spd-2-NG | Generated in this study; appears to be fully functional and rescues *Spd-2<sup>-/-</sup>* mutant. |
| Ubq-Spd-2-Cdk20-NG | Generated in this study; male sterile |
| Ubq-NG-Cnn-Cdk6 | Generated in this study; male sterile |
| Ubq-Spd-2 | Generated in this study |
| Ubq-Spd-2-Cdk20A | Generated in this study; male sterile |
| Ubq-Cnn | Generated in this study |
| Ubq-Cnn-Cdk6A | Generated in this study; male sterile |
| Ubq-Spd-2-mCherry | (Alvarez-Rodrigo *et al*, 2019) |
| Ubq-Jupiter-GFP | (Karpova *et al*, 2006)4/25/23 5:28:00 PM |

**Table 2:**
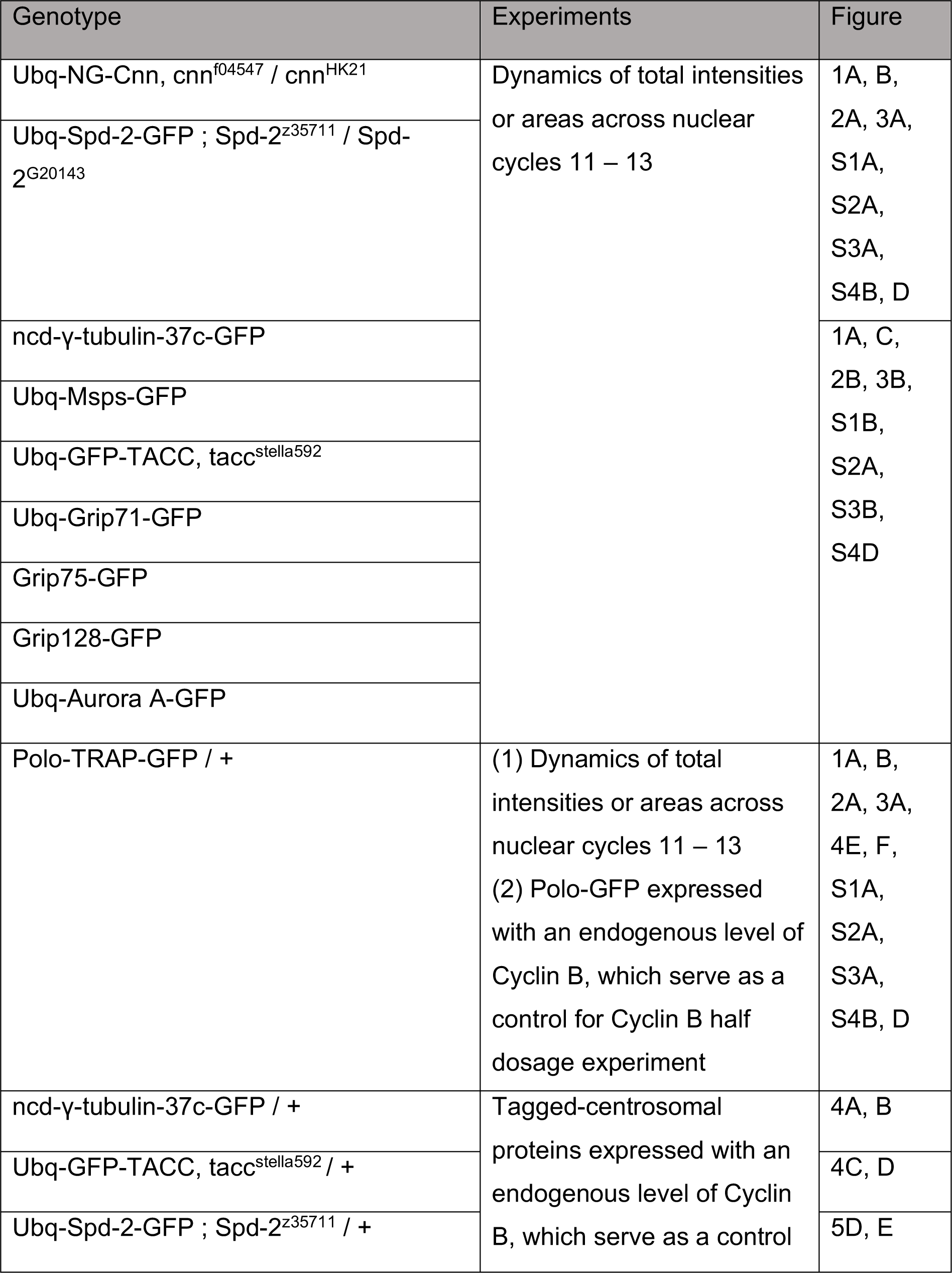
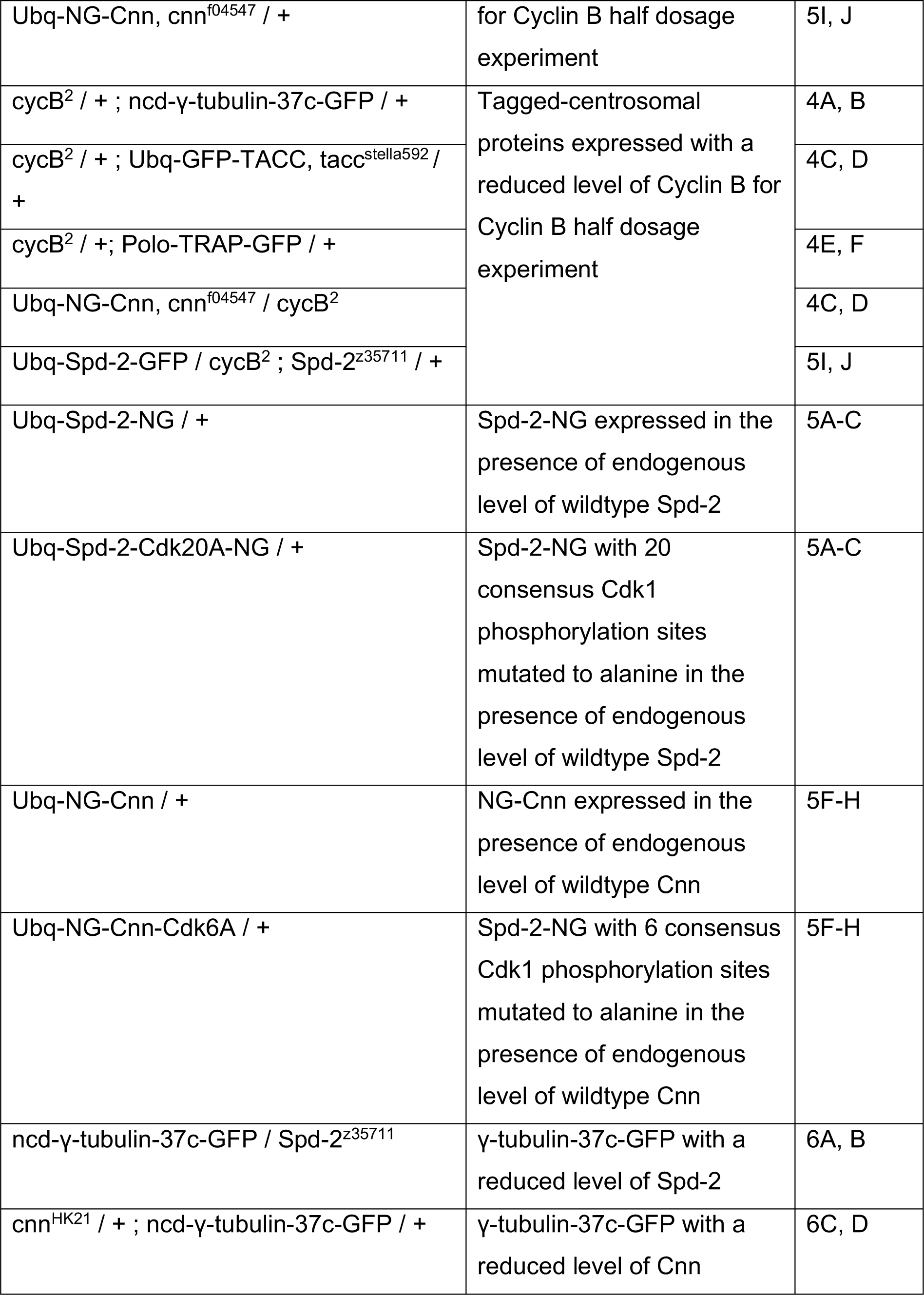
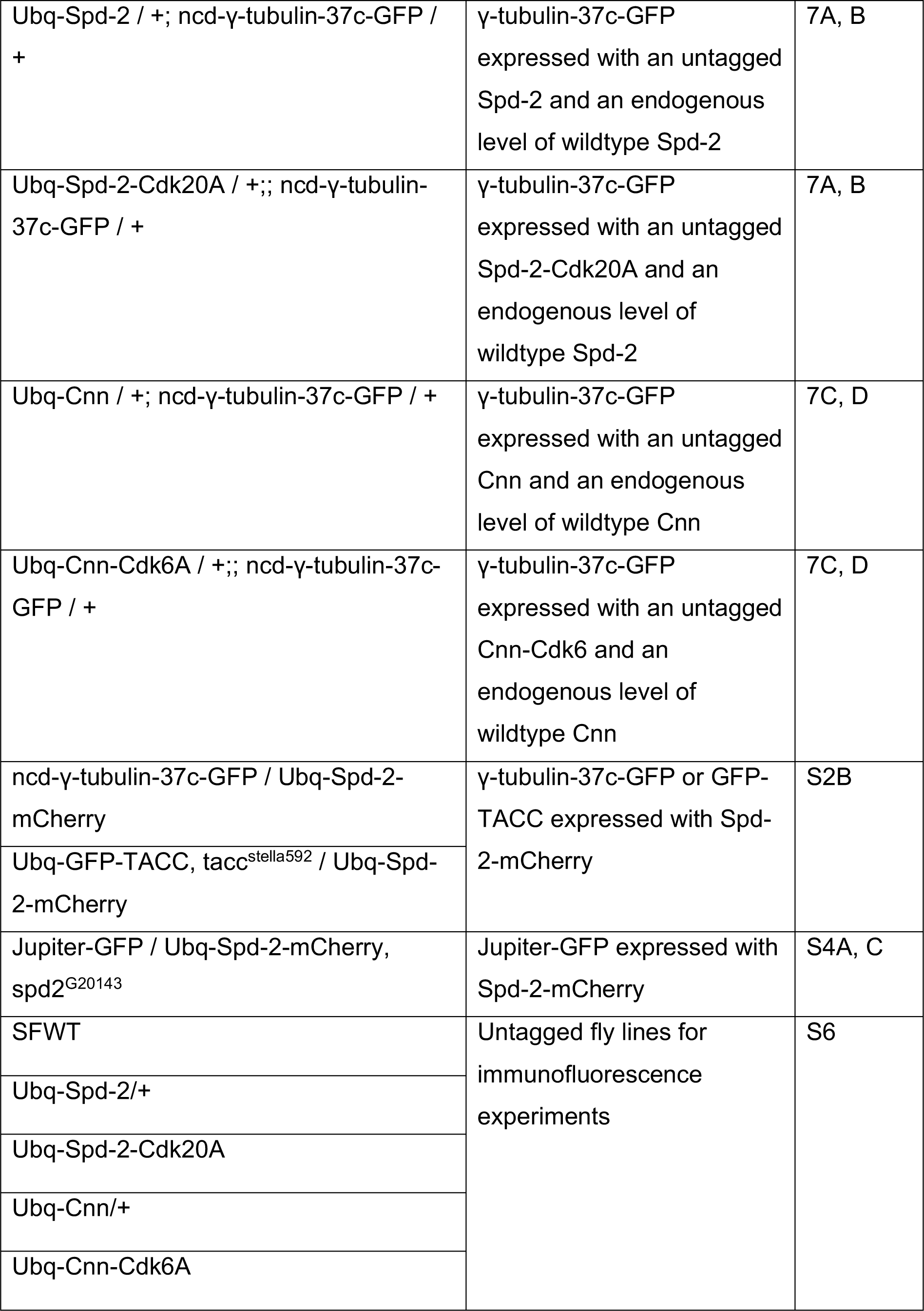
*Drosophila stocks* used in specific experiments.

### Transgenic fly line generation

Transgenic fly lines were generated via random P-element insertion (injected, mapped, and balanced by ‘The University of Cambridge Department of Genetics Fly Facility’). For transgene selection, the *w^+^* gene marker was included in the transformation vectors and constructs were injected into the *w*^1118^ genetic background.

To generate Spd-2-Cdk20A mutants, two different cDNA fragments encoding the following amino acid substitutions : S49A; T112A; S311A; T337A; S484A; T516A; S531A, S536A; T561A; S606A; S614A; S618A; S625A; S944A; S1021A;T1023A; S1065A; S1095A;S1102A;S1117A were synthesized by Genewiz and assembled with the PCR amplified backbone of pRNA-NG (CT) (Aydogan et al, 2020b) using HiFi Assembler (NEB, USA) to create Spd-2-Cdk20A-NG. This was recombined to create a pDONR vector and then a destination vector using Gateway technology (Thermo Fisher Scientific). For untagged Spd-2-Cdk20A, a stop codon was reintroduced to the C-terminus of the Spd-2-Cdk20A pDONR using Q5 Site Directed Mutagenesis (NEB, USA) and the resulting vector was recombined with a destination vector encoding no tag (Aydogan et al, 2018a), using Gateway technology. To generate Cnn-Cdk6A mutants, two different cDNA fragments (from the cnn-pA isoform, most highly expressed in embryos) encoding the following amino acid substitutions S64A; S91A; S364A; S1020A; S1057; S1067 were synthesized by Genewiz and assembled as described above. The resulting construct was recombined to create a pDONR vector and then a destination vector encoding mNG (NG-Cnn-Cdk6A) using Gateway technology. For untagged Cnn-Cdk6A, the Cnn-Cdk6A pDONR (described above) was recombined with a destination vector encoding no tag. Transgenic control lines expressing NG- or untagged-WT Spd-2 or WT Cnn were generated using the same DNA templates and methods but without mutagenesis.

#### Embryo collections

Embryos were collected from plates (40% cranberry-raspberry juice, 2% sucrose, and 1.8% agar) supplemented with fresh yeast suspension. For imaging experiments, embryos were collected for 1h at 25oC, and aged at 25oC for 45–60 min. Embryos were dechorionated by hand, mounted on a strip of glue on a 35-mm glass-bottom Petri dish with 14 mm micro-well (MatTek), and desiccated for 1 min at 25oC before covering with Voltalef grade H10S oil (Arkema).

#### Immunoblotting

Immunoblotting analysis to estimate protein expression level was performed as previously described (Aydogan, et al., 2018). The samples were lysed at 95°C for 10 min on a heat block, gently spun for 5 mins on a small lab bench centrifuge, and stored at −20°C. A total of 10 μl of the sample (which is the equivalent of 5 embryos for Cnn and 10 embryos for Spd-2) was loaded into each lane of a 3–8% Tris-Acetate pre-cast SDS-PAGE gel (Invitrogen, Thermo Fisher Scientific) and then transferred from the gel onto a nitrocellulose membrane (0.2 μm #162-0112; Bio-Rad) using a Bio-Rad Mini Trans-Blot system. For Western blotting, the membranes were incubated with blocking buffer (1× TBS + 4% milk powder + 0.1% Tween20) for 1h on an orbital shaker at 4°C overnight in blocking buffer with the primary antibody (see description below). The membranes were washed 3x with TBST (TBS + 0.1% Tween-20) and then incubated for 1h in blocking buffer with the secondary antibody (1:3,000 dilution, horseradish peroxidaseconjugated for chemiluminescence analysis). The membranes were washed 3x for 15 min with TBST buffer, before incubation for 1 min in HRPO substrate (Thermo Fisher Scientific SuperSignal West Femto Maximum Sensitivity Substrate, #34095) at a concentration that was empirically determined for each different protein and exposed to X-ray film for approximately 10 s to 2 mins. The following primary antibodies were used: rabbit anti-Spd-2 (1:500) [Lab stock #57] (Dix & Raff, 2007), rabbit anti-Cnn (1:1000) [Lab stock #37] (Lucas & Raff, 2007), and rabbit anti-GAGA factor (1:500) [Lab stock #144] (Raff *et al*, 1994). HRP-conjugated donkey anti-rabbit (NA934V lot:17876631, Cytiva Lifescience) secondary antibodies were used at 1:3000.

#### Spinning disk confocal microscopy

Images of embryos were acquired at 23°C using a PerkinElmer ERS spinning disk confocal system mounted on a Zeiss Axiovet 200M microscope using Volocity software (PerkinElmer). A 63X, 1.4NA oil objective was used for all acquisition. The oil objective was covered with an immersion oil (ImmersolT 518 F, Carl Zeiss) with a refractive index of 1.518 to minimize spherical aberration. The detector used was a charge-coupled device (CCD) camera (Orca ER, Hamamatsu Photonics, 15-bit), with a gain of 200 V. The system was equipped with 405nm, 488nm, 561nm, and 642 solid-state lasers (Oxxius S.A.). The microscope was operated using a Volocity software. All red/green fluorescently tagged samples were acquired using UltraVIEW ERS ‘Emission Discrimination’ setting. The emission filter of these images was set as followed: a green long-pass 520nm emission filter and a red long-pass 620nm emission filter. For dual channel imaging, the red channel was imaged before the green channel in every slice in a z-stacks. 0.5-μm z-sections were acquired, with the number of sections, time step, laser power, and exposure depending on the experiment.

#### Data analysis

Raw time-series images were imported into Fiji (Schindelin, et al, 2012) and corrected for photobleaching using the exponential decay function. Images were z-projected using the maximum intensity projection function, and the background was estimated and corrected using an uneven illumination background correction (Soille, 2004). Centrosomes were tracked using the TrackMate Plug-In in Fiji (Tinevez *et al*, 2016). A custom Python script was then used to threshold and extract the fluorescence intensities and areas of all tracked centrosomes as they changed over time in each individual embryo, as previously described (Wong *et al*, 2021). To extract the features of the Spd-2, Polo, γ-tubulin, Msps, TACC, Grip71, Grip75, Grip128 and Aurora A oscillations we measured the *initial intensity* of the centrosomes as they first separated in early S-phase and their *maximum intensity* as their levels peaked; the time between these points represented the *growth period*, while the *growth rate* was calculated as: (*maximum intensity* – *initial intensity*)/*growth period*. The total intensities of Cnn, which exhibits extensive centrosomal flaring, were extracted as described previously (Wong *et al*, 2021). The spindle length and mean intensity were analysed in embryos co-expressing the MT marker Jupiter-GFP (Karpova *et al*, 2006) and Spd-2-mCherry. The spindle length was calculated by averaging the distances between all pairs of centrosomes labelled by Spd-2-mCherry. The intra-centrosomal distance was subsequently used to constructed a box with as aspect ratio of (10 pixel-width x spindle-length-height), within which total intensities of Jupiter-GFP were averaged.

#### Analysis of spermatocytes

Testes were dissected from males of the appropriate genotype and were dissected, fixed and stained, as described previously (Roque, et al, 2012). The following antibodies were used: Sheep anti-Cnn (1:500; animal SKS027, (Cottee *et al*, 2013)), guinea pig anti-Asl (1:500; animal # SKC123; (Roque *et al*, 2012)), Rat anti-Spd-2 (1:500; animal SKR100, (Franz *et al*, 2013)), Alexa Fluor 488 nm-conjugated anti-rat IgG (1:500; Thermo Fisher Scientific; A21208), Alexa Fluor 594 nm-conjugated anti-sheep IgG (1:500; Thermo Fisher Scientific; A11016) and Alexa Fluor 633 nm-conjugated anti-guinea pig IgG (1:500; Thermo Fisher Scientific; A21105). Samples were mounted in Vectashield medium with DAPI (Vector Laboratories; H-1200). Meiotic cells were then scored blind for presence/absence of cytoplasmic Spd-2 and Cnn foci.

#### Statistical analysis

The details of statistical tests, sample size, and definition of the centre and dispersion are provided in individual Figure legends.

## Supplementary Figure Legends

**Figure S1.**
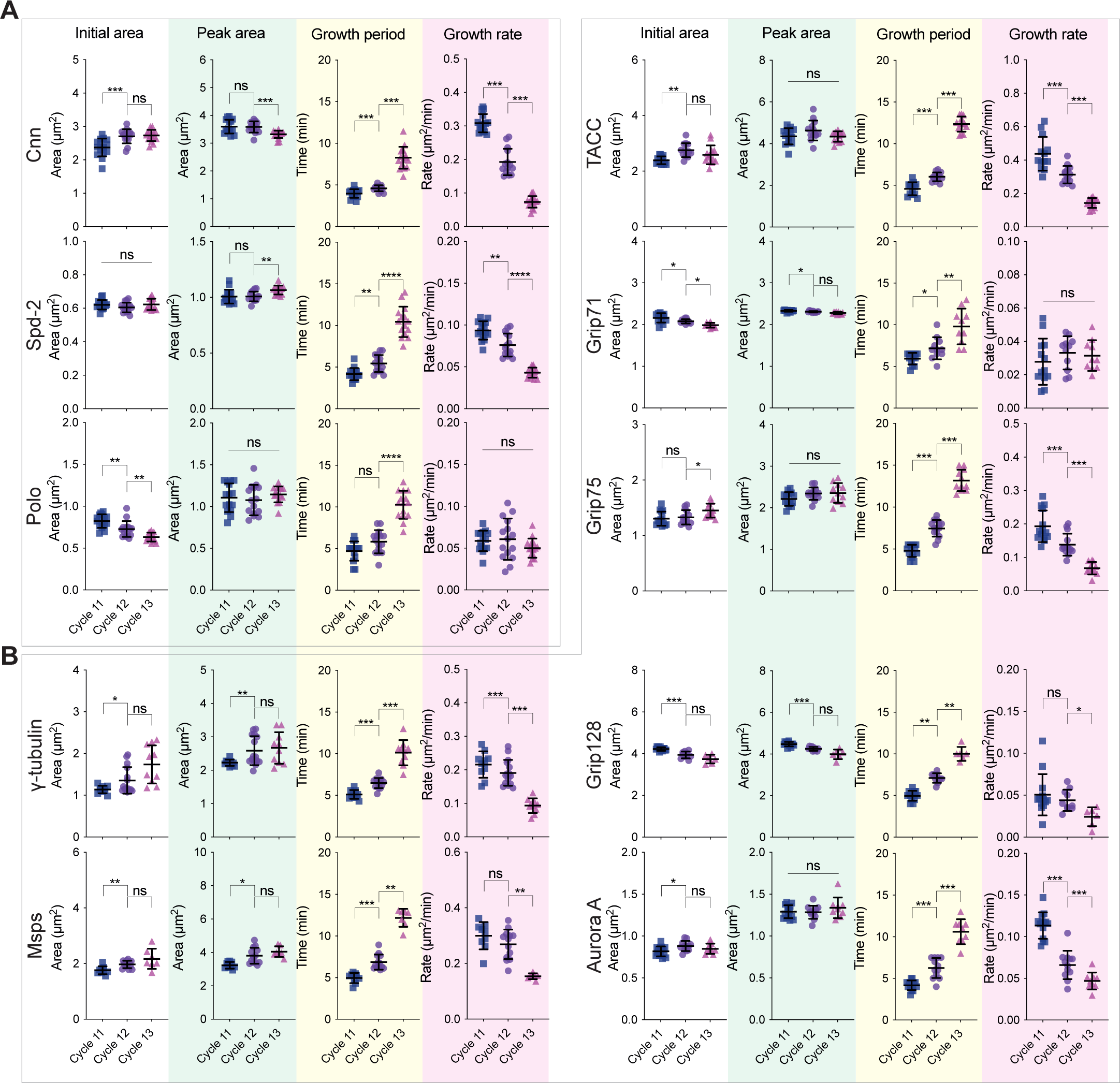
Analysis of centrosome growth kinetics measured by changes in centrosomal area during NC11, 12 and 13. **(A,B)** Graphs show how the mean centrosomal area (±SD of the data in each individual embryo shown in reduced opacity) of the PCM-scaffolding proteins Cnn, Spd-2 and Polo (A) and several PCM-client proteins (B) changes over time during NC11, 12, and 13. Note that these graphs were derived from the same embryos analysed in Figure 1. The length of each NC varies slightly from embryo to embryo, so all individual embryo tracks were aligned to NEB (t=0). The white parts of the graphs represent S-phase, and the grey parts represent mitosis. Note that the Grip71- and Grip128-fluorescent fusion proteins exhibited a very low signal-to-noise ratio, and this correlated with the proteins exhibiting very little change in their calculated centrosome-area during each nuclear cycle. N=7-15 embryos analysed at each nuclear cycle for each marker with a total of n=∼200-400, ∼400-800, or ∼600-1200 total centrosomes analysed at NC11, 12 and 13, respectively.

**Figure S2.**
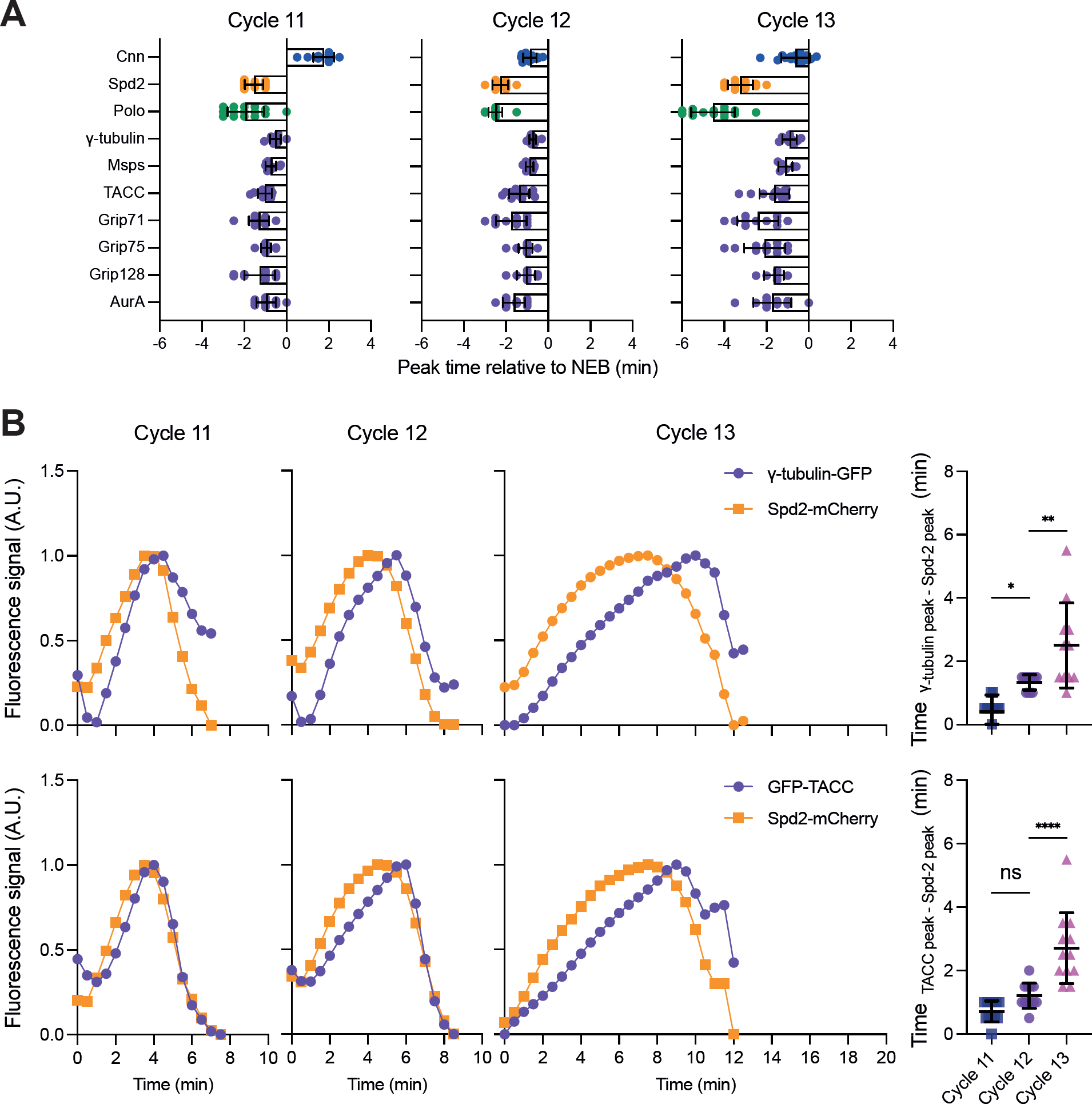
Analysis of the timing of the peak centrosomal fluorescence intensity of the various centrosomal proteins relative to NEB during NC11, 12 and 13. **(A)** Bar charts show the mean time (mins±SD) of the peak centrosomal fluorescence intensity relative to NEB (t=0) of the centrosomal proteins analysed here during NC11, 12, and 13 (calculated from the data shown in Figure 1). 10-15 embryos were analysed at each NC for each marker with a total of ∼200-300, ∼300-400, and ∼400-500 total centrosomes analysed at NC11, 12 and 13, respectively. **(B)** Graphs show the average centrosomal fluorescence intensity over time during nuclear cycles 11, 12 and 13 for embryos co-expressing Spd-2-mCherry (*orange*) with either γ-tubulin-GFP (*purple*, top graphs) or GFP-TACC (*purple*, bottom graphs). Fluorescence intensity was rescaled to between 0 and 1 in each cycle. The data from each embryo was aligned to centrosome separation at the start of S-phase (t=0). Accompanying dot plots compare the time difference between the Spd-2-mCherry peak and the γ-tubulin-GFP (top plot) or GFP-TACC (bottom plot) peak in each embryo. Data are presented as mean±SD. N=12 embryos analysed for each condition with a total of n=∼300, ∼500, or ∼900 total centrosomes analysed at NC11, 12 and 13, respectively. Gaussian distribution was tested using D’Agnostino and Pearson’s test. Statistical comparisons used a Kruskal-Wallis’s test (non-Gaussian-distributed). Multiple testing was performed using Dunn’s test.

**Figure S3.**
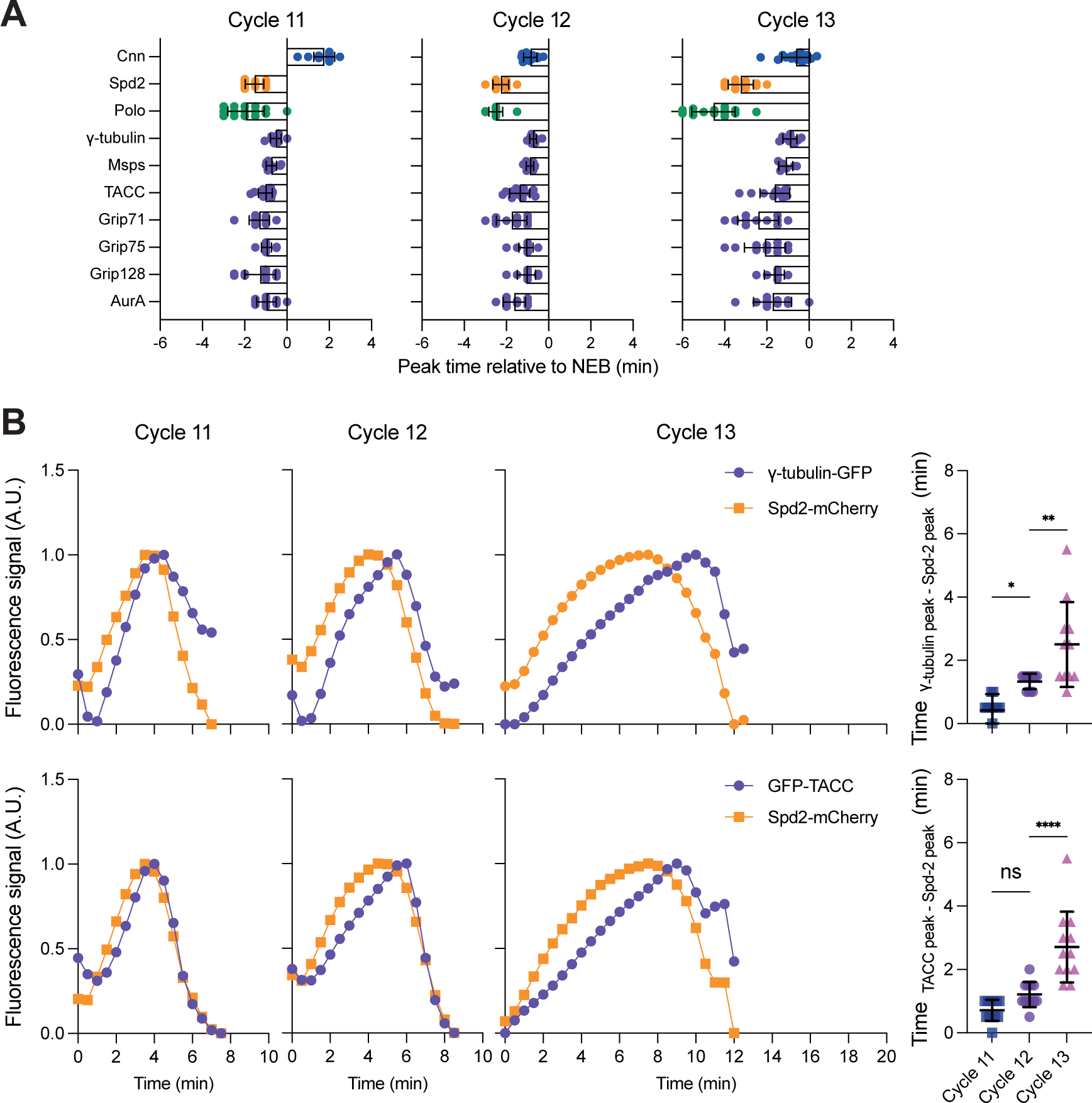
Analysis of centrosome growth parameters, measured by centrosome area, during NC11, 12 and 13. Scatter plots show the mean (±SD) initial fluorescent intensity (left graphs), peak fluorescent intensity (boxed in *green*), growth period (boxed in *yellow*), and growth rate (boxed in *pink*) in NC11, 12, and 13 for the PCM-scaffolding proteins **(A)** and PCM-client proteins **(B)**. Note that these graphs were derived from the same embryos analysed in Figure 1. Statistical comparisons used either an ordinary one-way ANOVA (Gaussian-distributed and variance-equal), a one-way Welch ANOVA (Gaussian-distributed and variance-unequal), or a Kruskal-Wallis’s test (non-Gaussian-distributed). If significant, multiple testing was performed using either Tukey-Kramer’s test (Gaussian-distributed and variance-equal), Games-Howell’s test (Gaussian-distributed and variance-unequal), or Mann-Whitney’s U test (non-Gaussian-distributed) (*: P<0.05, **: P<0.01, ***: P<0.001, ****: P<0.0001, ns: not significant). Gaussian distribution was tested using D’Agnostino and Pearson’s test. Variance homogeneity was tested using Levene W test.

**Figure S4.**
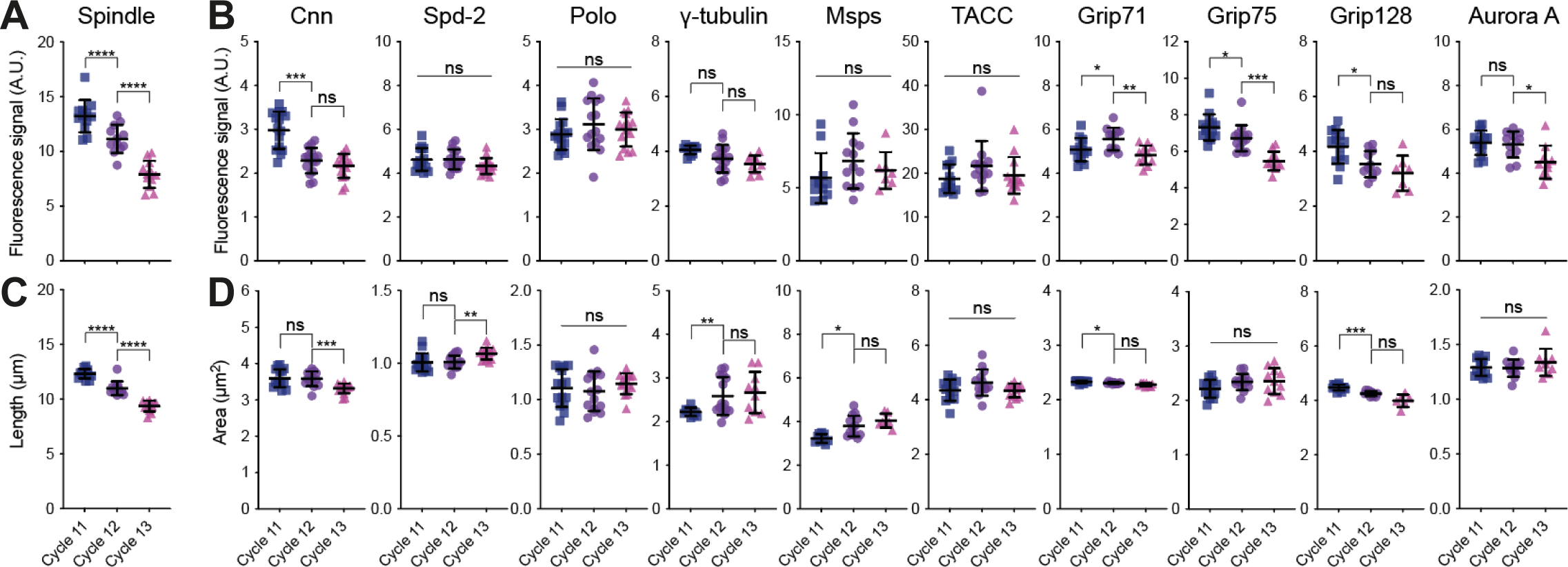
Comparison of how maximum spindle size and maximum centrosome size change during NC11, 12 and 13. **(A,B)** Scatter plots compare how the mean (±SD) fluorescence intensity of the metaphase mitotic spindle (A) and the mean (±SD) maximal fluorescence intensity of the various centrosome proteins analysed here (B) changes during NC11, 12 and 13. Note that the centrosome data shown in (B) is from the same peak intensity graphs shown in Figure 2B. **(C,D)** Scatter plots show how the mean length (±SD) of the metaphase mitotic spindle (C) and the mean maximal centrosome-area (±SD) of the various centrosome proteins analysed here (D) changes during NC11, 12 and 13. Note that the centrosome data shown in (B) is from the same peak area graphs shown in Figure S3. The metaphase spindles significantly reduce in size at each nuclear cycle (whether measured by spindle fluorescence intensity or spindle length), whereas centrosome size remains relatively constant (whether measured by centrosome fluorescence intensity or centrosome area). For the MT spindle measurements N=8-12 embryos were analysed at each nuclear cycle with a total of n=∼200-300, ∼400-500, or ∼600-800 total spindles analysed at NC11, 12 and 13, respectively. Statistical comparisons used either an ordinary one-way ANOVA (Gaussian-distributed and variance-equal), a one-way Welch ANOVA (Gaussian-distributed and variance-unequal), or a Kruskal-Wallis’s test (non-Gaussian-distributed). If significant, multiple testing was performed using either Tukey-Kramer’s test (Gaussian-distributed and variance-equal), Games-Howell’s test (Gaussian-distributed and variance-unequal), or Mann-Whitney’s U test (non-Gaussian-distributed) (*: P<0.05, **: P<0.01, ***: P<0.001, ****: P<0.0001, ns: not significant). Gaussian distribution was tested using D’Agnostino and Pearson’s test. Variance homogeneity was tested using Levene W test.

**Figure S5.**
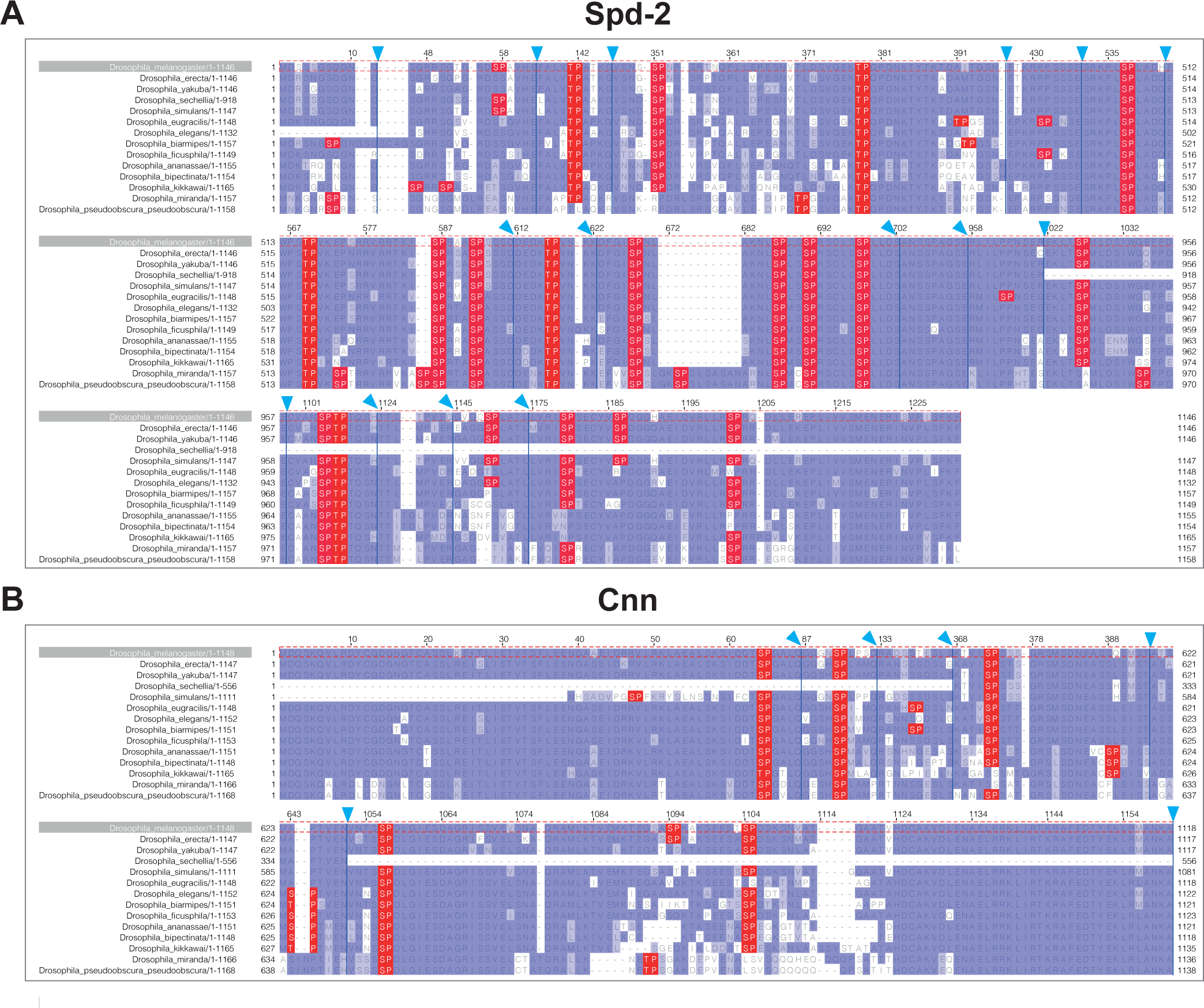
Analysis of potential Cdk/Cyclin phosphorylation sites (S/T-P motifs) in *D. melanogaster* Spd-2 and Cnn. Panels show multiple sequence alignments (MSA) of Spd-2 **(A)** and Cnn, highlighting the S/T-P motifs (*red*) present in the *D. melanogaster* proteins (top row of MSAs) that were mutated in this study and their levels of conservation in 13 other *Drosophila* species. The purple shading represents a BLOSUM62 conservation score (darker shading indicating higher conservation). The *blue arrows* above the MSA indicate the boundaries of regions of the MSA that are not shown here as they do not contain any S/T-P motifs mutated in this study. The sequences of the different *Drosophila* species were aligned using MUSCLE with default parameters and visualized using Jalview 2.11.

**Figure S6.**
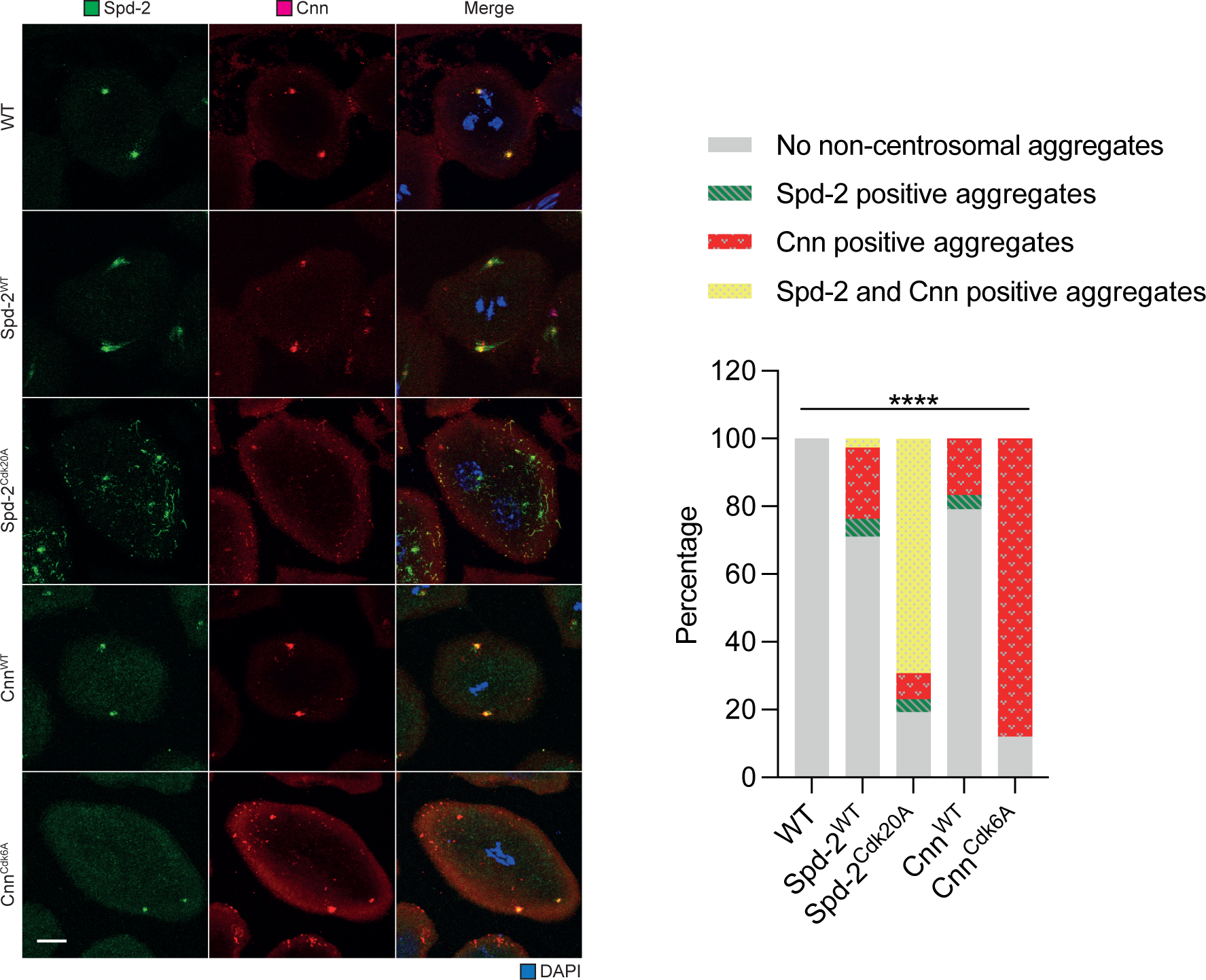
Analysis of meiosis in spermatocytes expressing Spd-2-Cdk20A-NG or NG-Cnn-Cdk6A. Images show examples of spermatocytes in meiosis I stained to reveal the distribution of Spd-2 (*green*), Cnn (*red*) or DNA (*blue*) from WT males, or males transgenically expressing either WT Spd-2, Spd-2-Cdk20A, WT Cnn or Cnn-Cdk6A. The spermatocytes expressing Spd-2-Cdk20A or Cnn-Cdk6A often contained prominent cytoplasmic aggregates; the Spd-2-Cdk20A aggregates often also recruit Cnn, but the Cnn aggregates do not detectably recruit Spd-2—in agreement with previous data showing that Spd-2 recruits Cnn to the mitotic PCM, but Cnn does not recruit Spd-2 (Conduit *et al*, 2014b). Note also that the transgenic expression of WT Spd-2 seems to lead to the recruitment of extra Spd-2 to the centrioles. Scale bar=10μm. (B) Bar chart quantifies the presence of these cytoplasmic aggregates in spermatocytes of the different genotypes— scored blind. N=10-50 spermatocytes per genotype. The statistical significance of the proportion change was calculated using the original number of spermatocytes by Chi-square test (****: P<0.0001)

## Notes

### Competing Interest Statement

The authors have declared no competing interest.

